# Short-chain mono-carboxylates as negative modulators of allosteric transitions in GLIC, and impact of a pre-β5 strand (Loop Ω) double mutation on crotonate, not butyrate effect

**DOI:** 10.1101/2023.03.14.530991

**Authors:** Catherine Van Renterghem, Ákos Nemecz, Karima Medjebeur, Pierre-Jean Corringer

## Abstract

The bacterial model GLIC remains one of the best known among pentameric ligand-gated ion channels (pLGICs), regarding their structure. GLIC is activated at low extracellular pH, but no agonist compound is known. Van Renterghem *et al*. (2023) showed that short-chain di-carboxylates potentiate GLIC activity, with strict dependence on two carboxylate binding pockets, previously characterized by crystallography (Sauguet *et al*., 2013, Fourati *et al*., 2015, 2020). An “in series” model was proposed, with compound binding at the inter-subunit pocket [homologous to the pLGICs orthotopic neutotransmitter binding site], and with involvement of the intra-subunit (or vestibular) pocket in coupling binding to gating.

Here we characterize saturated, mono-carboxylates as negative modulators of GLIC, as previously shown for crotonate (Alqazzaz *et al*., 2016). Butyrate and crotonate have indistinguishable properties regarding negative modulation of WT GLIC. However, a double mutation in the pre-β5 strand (Loop Ω) converts crotonate, as well as caffeate, but not butyrate, into positive modulators. We perform a mutational analysis of residue dependency in the two pockets, with the pre-β5 strand either intact or mutated. We propose that positive modulation requires stringent conditions, with integrity of both pockets, whereas negative modulation is less labile. The vestibular pocket may be involved as an accessory binding site leading to negative, but not positive modulation. We propose that the pre-β5 strand is involved in ligand-elicited modulation of GLIC gating, not in pHo-controlled gating. Possible involvement in Eukaryote pLGICs of regions corresponding to the vestibular pocket and the pre-β5 strand/Loop Ω is discussed.

**Key points summary:**

- Using the bacterial proton-activated receptor-channel GLIC, we identify a locus in the pre-β5 strand (Loop Ω) whose mutation inverses the effect of the mono-carboxylate crotonate from negative to positive modulation of the allosteric transitions, suggesting an involvement of the pre-β5 strand in coupling the extracellular orthotopic receptor to pore gating in this pentameric ligand-gated ion channel.
- As an extension to the previously proposed “in series” mechanism, we suggest that a orthotopic/orthosteric site – vestibular site – Loop Ω – β5-β6 “sandwich” - Pro-Loop/Cys-Loop series may be an essential component of orthotopic/orthosteric compound-elicited gating control in this pentameric ligand-gated ion channel, on top of which compounds targeting the vestibular site may provide modulation.

## Introduction

In human body, most pentameric ligand-gated ion channels (pLGICs) get activated by binding of neurotransmitters (or paracrine substances) to a major binding site, called the orthosteric site, or orthotopic site. This reference binding site, involved in physiological agonist effects, is located in the extracellular domain (ECD), at the interface between neighbour subunits, and accessible from the periphery of the pentamer. Many substances (toxins, toxic alcaloids, and chemistry products) are pharmacologically active on pLGICs (among which pesticides, convulsivants, drugs of abuse, and major clinically relevant substances). For many of them, the binding site is located in the transmembrane domain (TMD): either between TMD helices (propofol and several general anaesthetics, alcohols, barbiturates), or within the pore lumen (picrotoxinin, pumiliotoxin, ivermectine, lindane, etc.). Regarding agents active through the ECD, they very generally bind to the reference binding site (called orthotopic site in this report): either to the main orthotopic agonist sites (nicotine, alpha-bungarotoxin, etc.), or to an accessory orthotopic site in some heteromeric pLGICs (benzodiazepines on GABA_A_ receptors).

A new intra-subunit (intra-SU) binding pocket (also called vestibular pocket) was identified by crystallography in the ECD of several Prokaryote pLGICs. The vestibular pocket is adjacent to the inter-subunit (inter-SU) pocket (which is homologous to the reference pLGICs’ agonist site). It was identified in ELIC (Spurny *et al*., 2012), GLIC (Sauguet *et al*., 2013, Fourati *et al*., 2015, 2020; see also acetate in Nury *et al*., 2010), and sTeLIC (Hu *et al*., 2018). The question arises whether a homologous vestibular site may become a functional drug target in Eukaryote pLGICs, a question addressed by several authors (Hu *et al*., 2018, Brams *et al*., 2020).

Here we take advantage of the *Gloeobacter violaceus* ligand-gated ion channel (GLIC), very well characterized regarding structures, to analyse how the vestibule site may be involved in compound-elicited modulation of channel gating. In previously published GLIC-carboxylates co-crystal structures (Sauguet *et al*., 2013, Fourati *et al*., 2020), each compound tested was present in the co-crystal inter-SU (orthotopic) pocket. Some of them were also identified in the intra-SU (= vestibular) pocket (see Table 1), and no compound was found elsewhere than in the two pockets, therefore referred to as CBX-binding pockets. Van Renterghem *et al*. (2023) showed that integrity of the vestibular pocket is required for the modulation occurring by binding to GLIC inter-SU (orthotopic) site. Positive modulation by di-carboxylic acid/carboxylate (di-CBX) compounds (fumarate, succinate), and negative modulation by caffeate, showed an “all-or-none” pattern of residue-dependency: alanine replacement of a single residue in the inter-SU pocket (orthotopic site), or in the intra-SU pocket (vestibular site), abolished compound-elicited modulation.

**Table 1.**
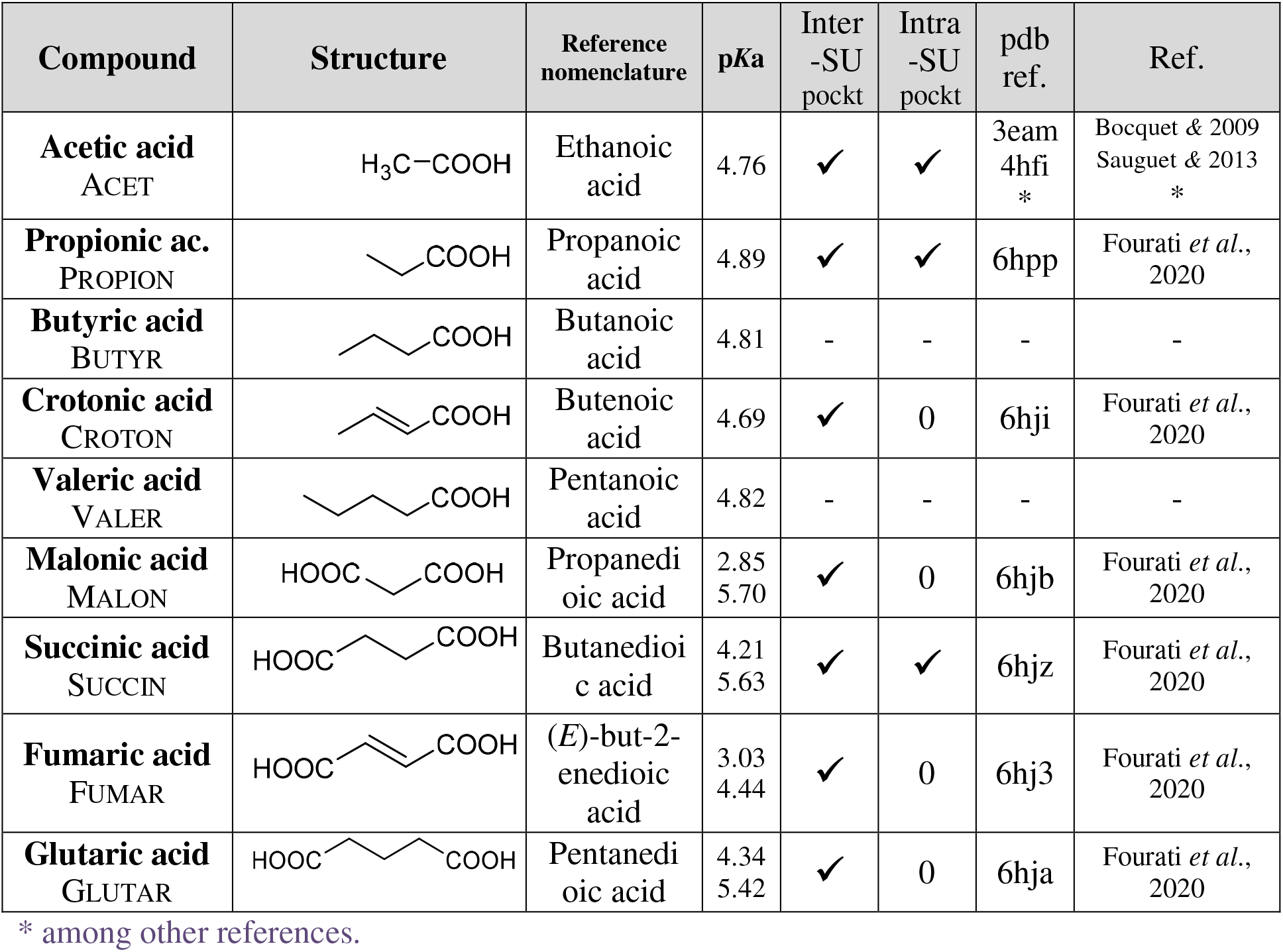
Mono-CBX and di-CBX compounds: names, structure, acidity constants, and presence in GLIC ECD carboxylate-binding pockets in published co-crystal structures. Acidity constants *K*a (given as p*K*a=-log(*K*a)) were obtained from the program Dozzzaqueux (10 mM at 25°C, with 170 mM NaCl).

In the present report, we use crotonic acid/crotonate (CROTON), previously identified as a GLIC inhibitor acting via GLIC ECD (Alqazzaz *et al*., 2016), and identify saturated, short-chain mono-CBX compounds (see Table 1) as negative modulators of the allosteric transitions in GLIC. We establish their “loose pattern” of residue dependency regarding the two CBX-binding pockets. Alanine substitution of Arg77, at the border between the two pockets, has a major impact on the ability of mono-CBXs to negatively modulate GLIC. Other substitutions in the CBX-binding pockets have either no impact or a relatively weak impact, leading us to propose that a (secondary) binding to the intra-SU site may modulate the influence of (primary) compound binding to the inter-SU site, and possibly mediate negative modulation, at least with a mutated orthotopic site.

In addition, we show that a double mutation in the pre-β5 strand, the pLGICs Loop Ω in the vestibule lumen, defined by Hu *et al*. (2018), inverses the effect of crotonate, from a strong negative modulation, into a strong positive modulation. This inversion occurs also with caffeate, but does not occur with butyrate, devoid of the double bond present in crotonate and caffeate. Negative modulation by crotonate (on WT GLIC) shows a “loose pattern” of residue dependency, whereas positive modulation by crotonate (on the pre-β5 variant) shows an “all-or-none” pattern of residue dependency. From our electrophysiology data, and previously published crystallographic data, we conclude that binding at the orthotopic site allows, through the vestibular region, and through the release of an Arg77-Asp88 inter-SU ion bridge, a pre-β5 strand motion involved in pore gating.

## Methods

### Electrophysiology

Electrophysiological methods were mostly as in Van Renterghem & Lazdunski (1994) and Van Renterghem *et al*., (2023), with heterologous expression, and using an RK-400 patch-clamp amplifier (or a two-electrode oocyte voltage-clamp amplifier), pClamp and a Digidata 1500 interface for control/acquisition, and Sigmaplot 11 for data analysis and Figures. We give here the main points and some differences.

### GLIC expression in tk-ts13 cells and *Xenopus laevis* oocytes

Apart for proton-activation curves (Fig. 6*B*), established using 2-electrode voltage-clamp from oocytes, electrophysiological data were obtained using whole-cell patch-clamp recording from tk-ts13 host cells, devoid of endogenous Acid Sensing Ion Channels (Van Renterghem *et al*., 2023), and here cultured without antibiotics. GFP-positive cells were used 1-2 days after calcium phosphate DNA transfection (tk-ts13 cells), or 1-3 days after nuclear injection (oocytes). A mixture of DNAs coding for GLIC and GFP in separate pMT3 vectors was used, with amounts of [2 + 0.2] µg per 35 mm dish, or [0.04 + 0.02] g/L in water for injections. GLIC residue numbering follows Protein Data Bank (PDB) entry 3EAM (Bocquet *et al*., 2009).

### Preparation of solutions

All solutions were prepared from the acid forms of the buffers/acido-basic compounds (BAPTA, HEPES, MES, CBX, etc.). Stock 100 mmol/L (mM) mono-CBX water solutions were prepared directly at pH 5 (NaOH/HCl), aliquoted and kept at -20 °C.

### Whole-cell patch-clamp recording

The intracellular pipette solution was composed of (in mM): CsCl 150, MgCl_2_ 1, HEPES 10, BAPTA 10, pH 7.3. The culture dish was washed and filled with the control extracellular solution (in mM): NaCl 165, MgCl_2_ 1, CaCl_2_ 1, MES 10, HEPES 6, pH 7.5 (NaOH to 9.5, HCl). Various solutions were applied by gravity, near the cell recorded, using either a multiway perfusion system converging to a single tip (50-100 µL/min, exchange time # 1 s) for ancient experiments (Figs 1*B*, 2, 3, and most of Figs. 4 & 5), or a computer-driven moving-head multichannel system (RSC-200, BioLogic), exchange time <30 ms in our conditions (Figs 1*A*, 7, 8 and part of Figs. 4 & 5). A 3 M KCl, 5 g/L agar bridge was used to connect the reference electrode to the dish solution. Transmembrane voltage was clamped most of the time to a constant value of -20 mV (occasionally -40 mV) [except during various non-detailed controls]. Electric current flowing through the membrane from extracellular to intracellular face (inward current) is counted negative, and represented downwards in the figures.

**Figure 1.**
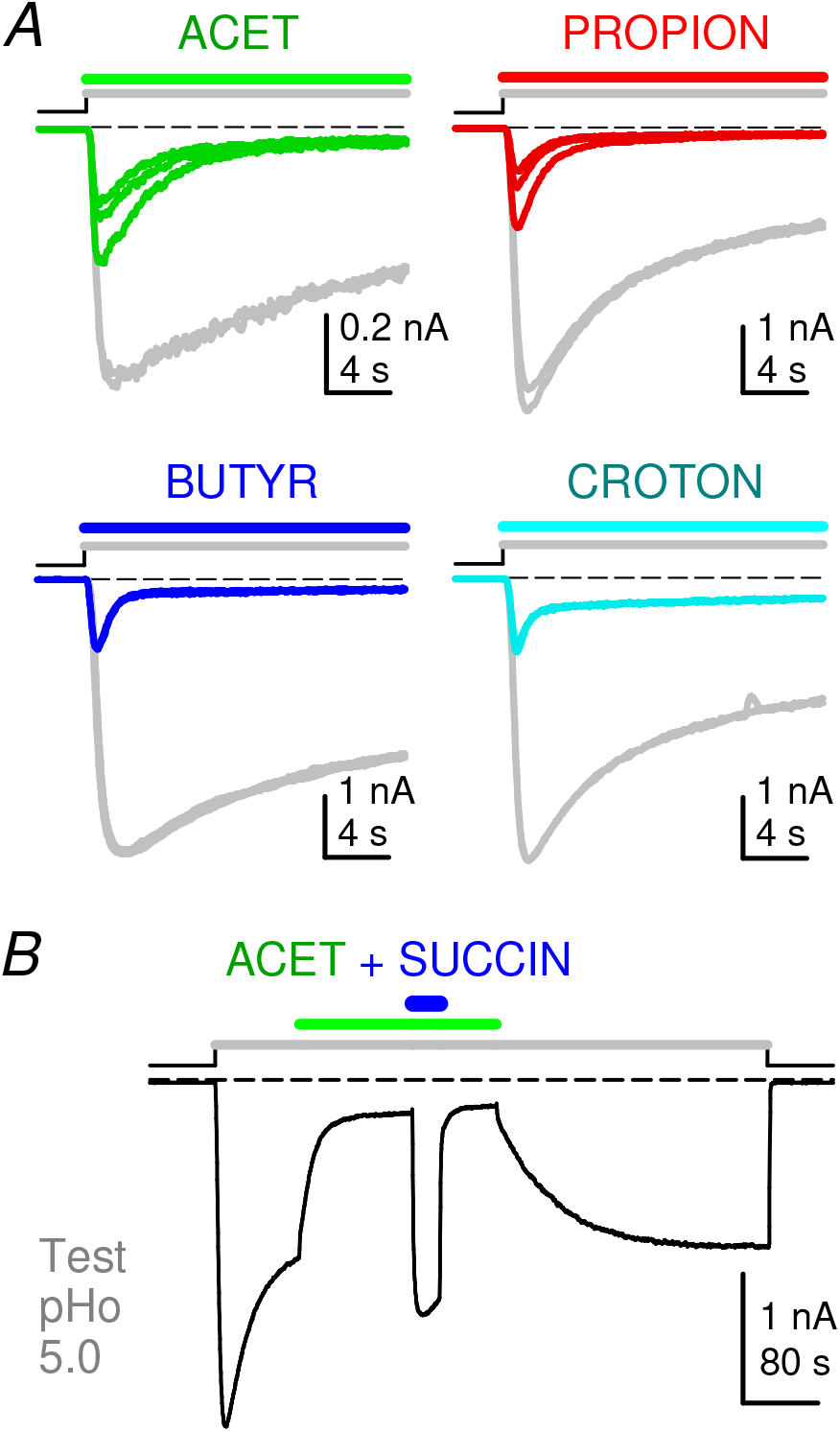
– Short chain mono-CBX compounds are negative modulators of allosteric transitions in GLIC. Traces of whole-cell voltage-clamp current recorded from tk-ts13 cells driven to express GLIC, with stimulations from control pHo 7.5. *A*, ***Sets of superimposed individual current traces (no averaging)*** recorded with stimulations at pHo 5.0 (20 s displayed), in the absence and in the presence of ACET, PROPION, BUTYR, and CROTON; see compounds in Table 1). Each set is from a different cell. Tests of 30 s duration were applied every 120 s, *i*. *e*. separated by 90 s wash times (no compound, pHo 7.5). Traces in *Grey* represent the last two stimulations at pHo 5.0 with no compound, and traces in *Colour* represent the subsequent first stimulations with a mono-CBX (1 mM at pHo 5.0; n = 3, 3, 2, 2 traces for ACET, PROPION, BUTYR, and CROTON respectively). *B*, After GLIC pre-stimulation at pHo 5.0, ACET (1 mM at pHo 5.0) was applied for 120 s, leading GLIC current to decrease to 18% of control (t = 13.6 s). SUCCIN (10 mM) was then added to ACET (pHo 5.0; 30 s), leading current to re-increase to 132 % of control (753% of current at 120 s ACET). Scale bars, Current: 0.2 nA (ACET) or 1 nA (others); Time: 4 s (*A*) or 80 s (*B*). Command potential: -40 mV (*A*) or -30 mV (*B*).

**Figure 2.**
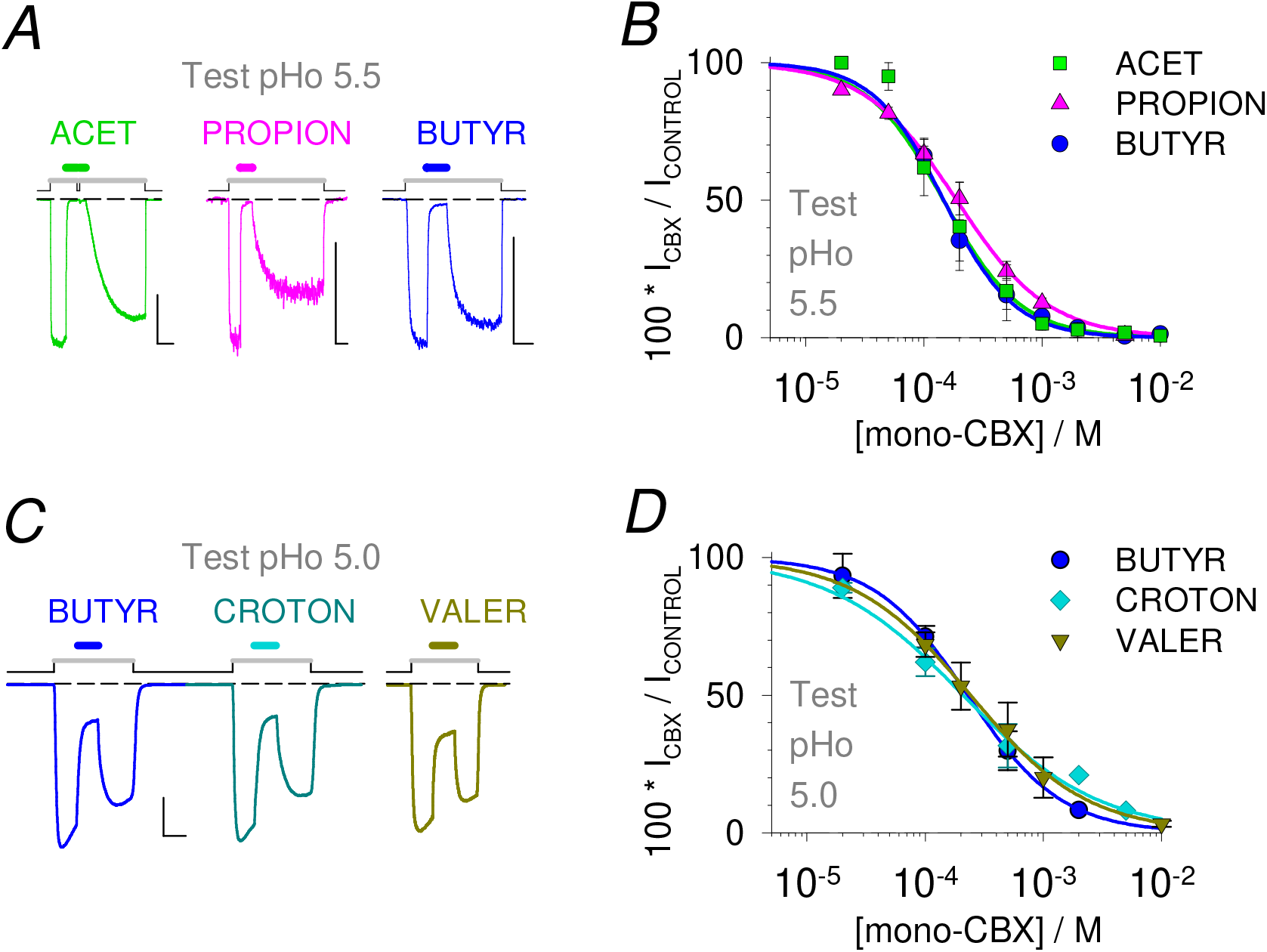
– No influence of carbon chain length or double bond on mono-CBX NAM effect. *A*, *B*, ***No impact of carbon-chain length***. *A*, Current traces showing activation of GLIC after switching pHo from 7.5 (*Black line* in protocol) to a lower test value (*Grey line* in protocol), here 5.5, followed by a reduction of GLIC current during application of mono-CBX solutions at pH 5.5 (*Colour lines*), and recovery from inhibition, followed by deactivation. From left to right are shown effects of the mono-CBX with two (ACET, 5 mM), three (PROPION, 5 mM) and four carbons (BUTYR, 2 mM). *B*, Corresponding plots of concentration-to-steady-state effect relations for ACET (constructed from 10 cells), PROPION, (n=4), and BUTYR (n=4), tested on GLIC current at pHo 5.5. Mean value ± SD. *C*, *D*, ***No impact of the double bond in CROTON vs BUTYR***. *C*, Current trace from one cell showing equal inhibitory effects of 0.5 mM BUTYR (saturated; *Blue line*), and of 0.5 mM CROTON (4-carbon, with a *trans* double bond; *Cyan line*), on WT GLIC activity at pHo 5.0. The trace with VALER (5-carbon saturated mono-CBX), also 0.5 mM at pHo 5.0, is from a different cell. *D*, Corresponding concentration-to-effect plots, showing equal IC_50_s for BUTYR (*Blue Circles*; n=5) and CROTON (*Cyan Diamonds*; n=4), and for VALER (*Dark yellow Triangles*; n=4), at pHo 5.0. Mean value ± SD. Scale bars, Current: 0.1 nA (PROPION) or 0.5 nA (others); Time: 60 s (ACET, PROPION) or 20 s (others).

**Figure 3.**
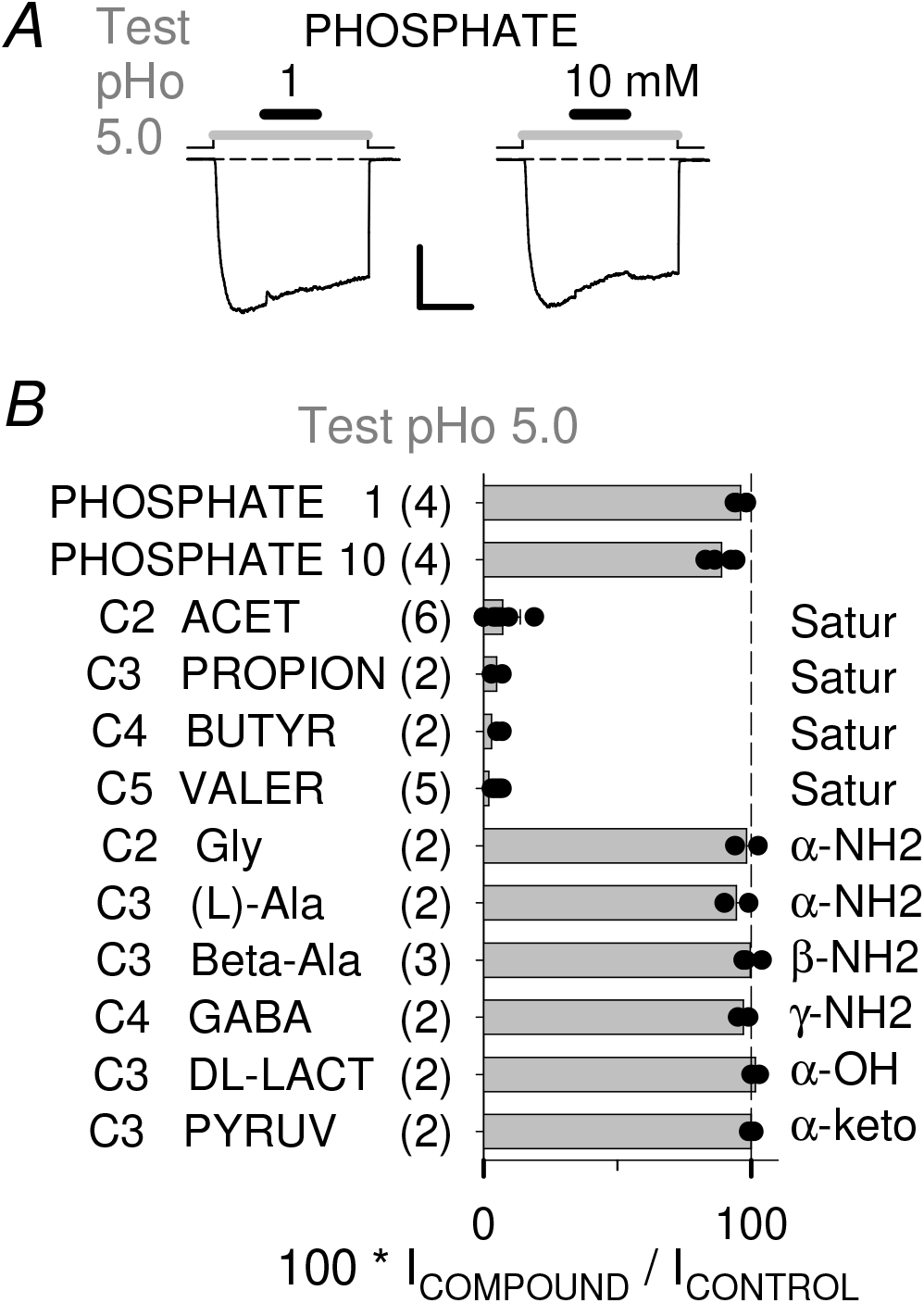
– Compound selectivity of the mono-CBX NAM effect on GLIC. *A*, Representative current traces for phosphate tests (1 and 10 mM at pHo 5.0). Scale bars, Current: 0.4 nA, Time: 20 s. *B*, Bar graph of current in the presence of a compound (in % of control pHo 5.0-elicited current; protocol with pre-stimulation) for phosphate (1 and 10 mM; in 0 Ca solution), mono-CBX (1 mM) and mono-CBX based compounds with amino-, hydroxy-or keto-groups (5-10 mM). Each category name also indicates the number of carbons (C2-C5), and the number of cells tested (in brackets). Mean value, with individual cells data points superimposed, and ± SD if n=3 or greater.

**Figure 4.**
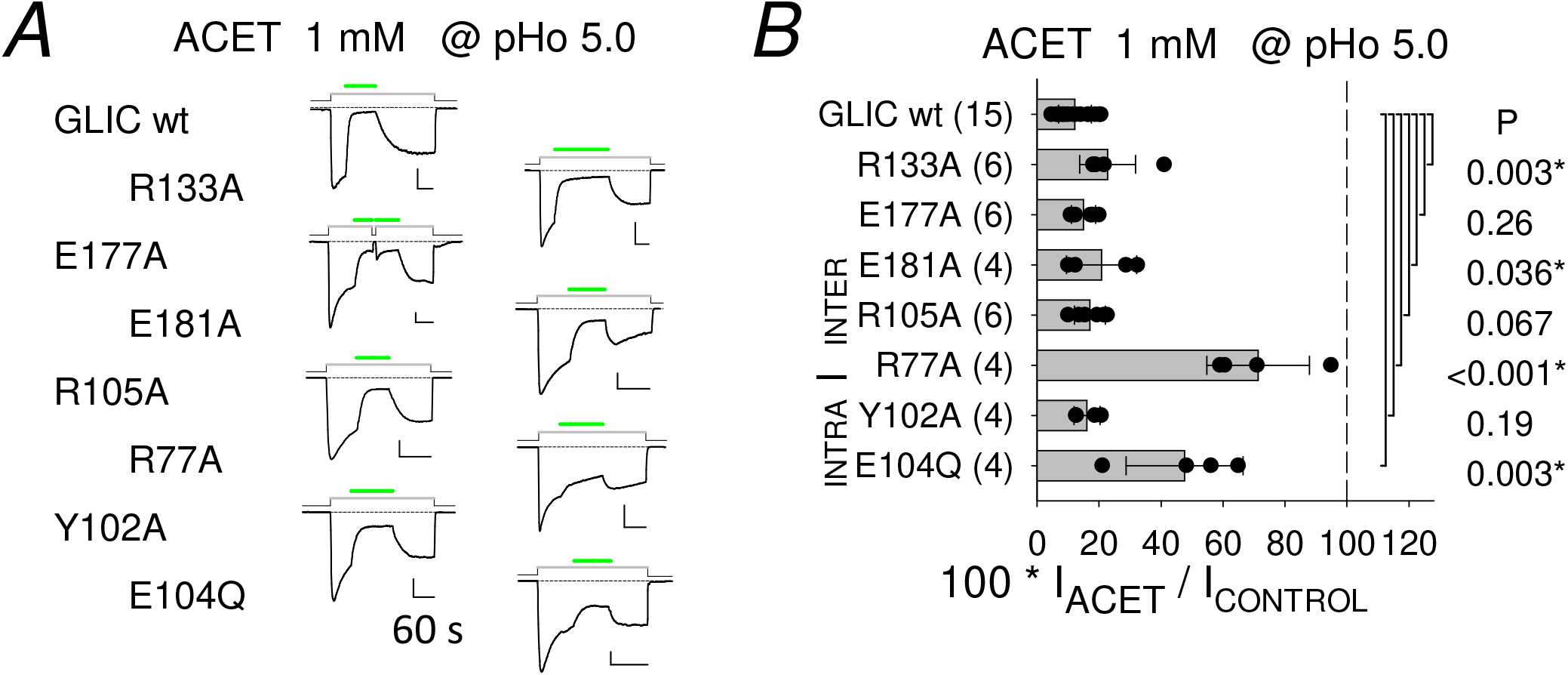
– CBX binding pockets single mutations: impact on ACET NAM effect. *A*, Representative current traces illustrating ACET tests (1 mM at pHo 5.0) on WT GLIC and single mutation GLIC variants, as indicated left to the traces. Scale bars, Current: 0.2 nA; Time: 60 s. *B*, Bar graphs of current measured after 60 s in the presence of 1 mM ACET (in % of control at pHo 5.0) on WT and single mutation GLIC variants, as indicated. Residue belonging to the inter-or intra-SU CBX-binding pocket is indicated, as well as the border/pivot Arg77 (*Bar*). Individual data points are superimposed to bars indicating mean ± SD, for each sample of cells tested, with the number of cells tested given in brackets. Student T-test P value is indicated for each mutant to WT pair of samples.

**Figure 5.**
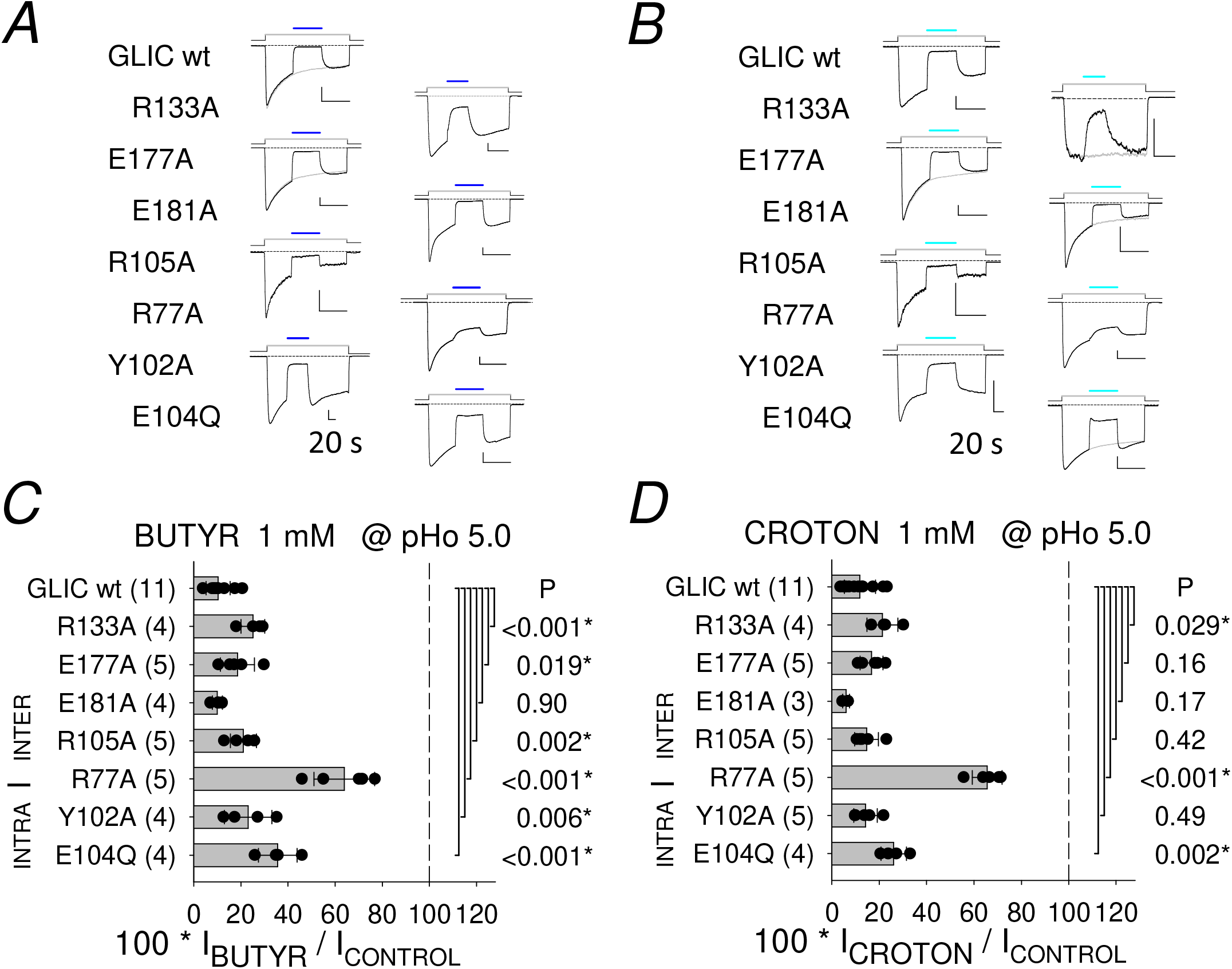
– CBX binding pockets single mutations: impact on BUTYR and CROTON NAM effects. *A*, *B*, Representative current traces illustrating BUTYR (A) or CROTON (*B*) tests (1 mM at pHo 5.0), in the pre-stimulation protocol, on WT GLIC and single mutation GLIC variants, as indicated left to the traces. Scale bars: Current: 0.1 nA (BUTYR R105A; CROTON R105A and R133A), or 0.4 nA (others); Time: 20 s. *C*, *D*, Bar graphs of current in the presence of 1 mM BUTYR (*C*) or CROTON (*D*), (in % of control at pHo 5.0), on WT and single mutation GLIC variants, as indicated. Student T-test P value is indicated for each mutant to WT pair of samples.

### Two-electrode voltage-clamp from oocytes

Whole-oocyte voltage-clamp was performed using two 3 M KCl-filled intracellular pipettes containing Ag/AgCl electrodes, and two extracellular Ag/AgCl pellets connected to the bath using two separate agar bridges. The control extracellular solution was composed of (in mM): NaCl 100, KCl 3, MgCl_2_ 1, CaCl_2_ 1, MES 10, pH 8.0, in order to keep the conditions used by Nemecz *et al*. (2017) and Van Renterghem *et al*. (2023), although a different set-up was used in the present study, with an OC-725C amplifier (Warner Instruments), and manual fast chamber perfusion with an Omnifit solution exchange system.

### Pharmacology and binding sites

As in Van Renterghem *et al*. (2023), we use the words “ortho*topic*”/”allo*topic*” to characterize the *location* of a binding site, and “allo*steric*” to comment *3D conformational* changes. Therefore (1A/) we use “orthotopic site” [in replacement of the more usual “orthosteric site”] to designate the reference agonist site in pLGICs [the neurotransmitter binding site], as well as the homologous location in GLIC, [including Arg105, Arg133 (Loop B), Asp177 (Loop C), Asp181]. 1B/ According to Sauguet *et al*. (2013) and Fourati *et al*. (2020), the inter-subunit CBX binding pocket in GLIC, [including Arg77 (Loop A), Arg105 (Loop E), Asp181 (Loop C), and, for the di-CBXs, Asn152 (Loop F)], is overlapping with the conserved orthotopic site, but situated a bit more deeply, and accessible from the periphery of the pentamer through an entrance which is part of the orthotopic site (Arg133, Asp177). 2/ The intra-subunit CBX binding pocket in GLIC [including Arg77 in the apo-GLIC, and Arg85 (Pre-β5), Tyr102 (β6), Glu104 (β6)], accessible from the vestibule *lumen*, corresponds exactly to the “vestibular” pocket in ELIC and other pLGICs, and constitutes an “allotopic” binding site, involved in the control of allosteric transitions. When discussing (1/) *vs* (2/), we may occasionally use “orthotopic” to designate the ensemble (1B+1A) (inter-SU CBX binding pocket + the conserved site), as opposed to the “allotopic”, intra-SU (= vestibular) pocket (2/). In WT GLIC activated at low pHo, keeping as a reference the pLGICs reference binding site, fumarate is an orthotopic PAM, crotonate an orthotopic NAM, propofol an allotopic NAM, extracellular proton an allotopic agonist. The same vocabulary is perfectly consistent regarding Eukaryote pLGICs (neurotransmitters are orthotopic PAMs with agonist property, *i*.*e*. orthotopic agonists, propofol an allotopic PAM or NAM, etc.).

In most experiments, a protocol with pre-stimulation was used (Figs. 1*B*, 2, 3, 4, 5, 7, 8). Compound application started after 60 s of GLIC stimulation at low extracellular pH (low pHo) in experiments with acetate 0.5-10 mM (Figs. 1*B*, 2*AB*, 4), propionate (Fig. 2*AB*), and compounds in Fig. 3 except for phosphate (20 s) (and after up to 180 s for lower acetate concentrations). For compounds with faster effect (butyrate, crotonate, valerate), pre-stimulation time was 20 s (Figs. 2, 7 and about two third of cells in Fig. 5), and 10 s in Fig. 8. Fig. 5 includes for each construct about one third of ancient experiments using 60 s pre-stimulation, then 20 s- and 60 s-data have been pooled in analysis.

Positive/negative modulation was evaluated as current in the presence of the compound, in percent of the control GLIC current value measured immediately before compound application (with a correction for GLIC current decay only for small inhibitions). The numerical values which are given in text (mean ± SD), used for statistical evaluations, and plotted in graphs are measures of this parameter (100*I_CBX_/I_CONTROL_) [occasionally called “PAM *ratio* (%)”, and comprised between 0 and 100 in NAM effects].

Fit of dose-response curves. For proton activation curves, the peak value (Ipk[pHo]; inward current value at maximal absolute value) within 30 s was considered (y), and a plot against H^+^ activity values (x) was fitted with a 3-parameter Hill equation: y = y_max_ *{1 / [1 + (EC_50_^n_H_ / x^n_H_)]}, giving I_max_, EC_50_, and pH_50_ = -log(EC_50_). For potentiation/inhibition, data for percent of current in the presence of the compound (y = 100*I_CBX_/I_CONTROL_) to compound concentration values (x), [which are presented in the Figures in log-scale x-axis display], were fitted using a 4-parameter Hill equation (potentiation: y = 100 + (y_max_-100) *{1 / [1 + (EC_50_^n_H_ / x^n_H_)]}), or a derived sigmoidal function (inhibition: y = y_max_ *{1 - 1 / [1 + (IC_50_^n_H_ / x^n_H_)])}), giving y_max_, EC_50_ or IC_50_, and an empirical slope parameter, noted n_H_, defined so that its values are positive. For most compounds are presented: a fit of the mean of data (Graphs), and the mean ± standard deviation (SD) of data from individual cells (values in text).

### Statistics

In the mutational analysis, the sample of (100*I_CBX_/I_CONTROL_) values for each mutant was compared to the sample for WT GLIC (Figs. 4, 5, and part of Fig. 7), or the sample for GLIC D86A-D88A (Fig. 8, and part of Fig. 7), using a Student’s T-test. The probability (P) for test and reference samples coming from a single normally distributed population is indicated near each mutant data. The *criterium* used in text for a significant impact of a mutation was: P < 0.05.

## Results

### Part 1 – Mono-CBX compounds NAM effect on WT GLIC

#### 1-1 / Short chain mono-carboxylates elicit a negative modulation of allosteric transitions in GLIC

A *direct protocol*, where a pHo-jump (from control pHo 7.5) and compound addition are applied simultaneously, was first used to examine the influence of saturated, short chain mono-CBX compounds on low pHo-induced GLIC current (Fig. 1*A*).

Using pHo 5.0 (corresponding approximately to pEC_50_ [or pHo_50_] on GLIC), low pHo induced GLIC current decay is relatively slow (see control traces, in Grey, in Fig. 1*A*: more than a half of peak current value remaining after 20s), in comparison with the decay of EC_50_ whole-cell patch-clamp currents from most mammalian pLGICs. If a new stimulation includes acetic acid /acetate (ACET; 1 mM at pH 5.0), added to the [MES + HEPES] – buffered solution (Fig. 1*A*, *Upper Left* trace), a decreased peak current value, and a faster current decay (than in the absence of ACET) are observed, revealing an inhibitory effect of ACET.

The 3- and 4-carbon saturated mono-CBX compounds, propionic acid/propionate (PROPION), and butyric acid/butyrate (BUTYR; both 1 mM at pHo 5.0), also elicited a reduced peak current and faster current decay. Alqazzaz *et al*., (2016), characterized the 4-carbon mono-CBX with a *trans* double bond, crotonic acid/crotonate (CROTON) as a GLIC inhibitor at pH 5.5. CROTON was also an inhibitor at pHo 5.0 (Fig. 1*A*).

For the first CBX application in each set (Fig. 1*A*), the *ratio* of peak current values in the presence and in the absence of compound (Ipk[CBX@5.0] / Ipk[pHo 5.0]) was evaluated. Ratios were 0.55 (ACET), 0.37 (PROPION), 0.24 (BUTYR), and 0.26 (CROTON), for the cells in Fig. 1*A*, showing that a fast inhibition component occurs during GLIC activation in the conditions used. The current (absolute value) time course was fitted with a Hodgkin & Huxley function *m*^4^*h* (*m*, a rising exponential function of time; *h*, a decay exponential), giving for curent decay a time constant of 3.3 s (ACET), 1.5 s (PROPION), 0.54 s (BUTYR), and 0.58 s (CROTON) on these cells.

A *protocol with prestimulation* is used in Fig. 1*B*: GLIC is pre-stimulated at low pHo (here 5.0) before addition of a compound (here ACET 1 mM at pHo 5.0). The 4-carbon di-CBX compound succinic acid/succinate (SUCCIN) was previously characterized as a positive modulator of the allosteric transitions (PAM) on GLIC activated at pHo 5.0 (Van Renterghem *et al*., 2023). Applied on a cell after reaching the plateau of inhibition by ACET (1 mM at pHo 5.0, 50 s, to 6.6 % of control on the cell commented), SUCCIN (10 mM at pHo 5.0, without ACET), produced a fast, reversible, re-increase of GLIC current to 143 % of the control current recorded immediately before ACET (representing 2170 % of the current recorded in the presence of ACET). A similar pattern was observed if, at the plateau of ACET inhibition, SUCCIN (10 mM) was co-applied with ACET (1 mM; pH 5.0; Fig. 1*B*): the PAM SUCCIN overcame ACET inhibitory effect. On 3 cells, the peak SUCCIN current values (co-application; after 90, 120 & 180 s ACET 1mM, reaching 5.5, 17.2, 11.3 % of control current at pHo 5.0) were: 28, 132 (Fig. 1*B*), & 44 % of the control current recorded immediately before ACET (representing 505, 753 (Fig. 1*B*), & 392 % of ACET current). Despite variability in its quantitative result, this functional competition experiment shows that inhibition by ACET does not occur through an inhibition of permeation, but occurs through a negative modulation of the receptor-channel gating transitions.

#### 1-2 / No influence of carbon-chain length or double bond on mono-CBX NAM effect on WT GLIC

A protocol with pre-stimulation at low pHo was chosen to further characterize mono-CBX NAM effects on GLIC, as illustrated in Fig. 2*A* for ACET (5 mM), PROPION (5 mM) and BUTYR (2 mM), at pHo 5.5. The protocol with pre-stimulation also shows that reversibility occurs after compound wash-out at low pHo. The relation of ACET concentration to minimum inward current value (in percent of control), is plotted in Fig. 2*B*. A sigmoid fit of the mean value (established using 10 cells) against concentration indicates an IC_50_ of 136 µM and a slope parameter equal to 1.6. Fitting the concentration to current (% value) relations from individual cells indicated an IC_50_ of 128 (± 55, *n* = 5) µM for ACET at pHo 5.5. With 3- and 4-carbon compounds, the IC_50_ and slope parameter values obtained from individual cells were for PROPION 204 (± 51, *n*=4) µM and 1.1 (± 0.1, n=4), and for BUTYR 150 (± 23, *n*=4) µM and 1.5 (± 0.2, n=4).

Changing the test pHo value from 5.5 to 5.0 had a significant but weak influence on BUTYR effect (Fig. 2*CD*), with an IC_50_ value of 219 (± 31, n=4; P=0.012) µM at pHo 5.0 *vs* pHo 5.5. The 5-carbon mono-CBX compound, valeric acid/valerate (VALER) showed an inhibitory effect, with an IC_50_ of 289 (± 116, n=4). CROTON NAM effect at pHo 5.0 (Fig. 2*CD*) was indistinguishable from BUTYR effect at the same pHo, with an IC_50_ value of 191 (± 63, *n* = 4; P=0.46) µM. Isocrotonic-acid/isocrotonate, the 4-carbon mono-CBX with a double bond in the *cis* configuration, is known to be unstable due to spontaneous *cis* to *trans* isomerisation at low pH. Having no simple tool to evaluate the proportion of *cis* compound, we decided not to test isocrotonate.

This data shows that mono-CBX compounds negatively modulate GLIC with no impact of the carbon chain length, no impact of a *trans* double bond in the 4-carbon compound, and no major influence of the test pHo value between 5.5 and 5.0. Using a pipette solution at pH 7.3, we found no condition in which ACET, PROPION, BUTYR or CROTON would produce a potentiation of WT GLIC current, no evidence for a biphasic effect according to concentration or pHo: only inhibitory effects were observed with mono-CBX compounds on WT GLIC.

#### 1-3 / Selectivity of GLIC for mono-carboxylates

A protein crystallization liquor buffered with phosphate, instead of acetate or another CBX, was used by Fourati *et al*. (2015) to obtain an Apo-GLIC crystal structure (PDB reference 4qh5), and no phosphate ions were identified in the structure. As an echo, we performed functional tests of phosphate on GLIC activity (recorded in a solution identical to our extracellular solution except that CaCl_2_ was omitted): as shown with current traces (Fig. 3*A*), phosphate (1 and 10 mM) had no effect on GLIC activity at pH 5.0 (n = 4 cells with two concentrations each; Fig. 3*B*).

Tested at 1 mM with pre-stimulation, ACET and PROPION were active at pHo 5.0 (Fig. 3*B*), as seen for BUTYR and VALER (Fig. 2*CD*; Fig. 3*B*). Neighbour compounds as well (5 or 10 mM) were tested with pre-stimulation at pHo 5.0 (Fig. 3): the amino-derivatives glycine (C2 as ACET) and (L)-alanine (C3 as PROPION), β-alanine (β-amino propionate) and γ-amino butyrate (GABA), as well as the α-hydroxy-derivative DL-lactate (C3 as PROPION) and α-keto-derivative pyruvate (C3 as PROPION). All these compounds had no effect on GLIC current (2-5 cells each), showing a strong chemical selectivity for the short chain mono-CBXs NAM effect on GLIC.

#### 1-4 / Functional relevance of GLIC CBX binding pockets for negative modulation by ACET, BUTYR and CROTON: mutational analysis (pHo 5.0)

The inter-SU and intra-SU CBX-binding pockets revealed in the GLIC protein by crystallographic structures (Sauguet *et al*., 2013, Fourati *et al*., 2020), and the overlapping pLGICs orthotopic site region, were previously evaluated (at pHo 5.0) for their involvement in the PAM effects of SUCCIN and fumaric acid/fumarate (FUMAR), and the NAM effect of caffeic acid/caffeate (CAFFE). Van Renterghem *et al*. (2023) showed that a mutation in anyone of these locations has an “all-or-none” impact, with suppression of both the 4-carbon di-CBX PAM effects, and (except for E181A) the CAFFE NAM effect. Here we used the same single mutants to test (at pHo 5.0) the functional relevance of these binding *loci* in the 4-carbon mono-CBX NAM effects and ACET effect. We show that most mutations have much less impact on the mono-CBX NAM effects, than on di-CBX and CAFFE effects.

ACET (Fig. 4) and the 4-carbon compounds BUTYR and CROTON (Fig. 5) were tested on the CBX-pockets single mutants, at 1 mM with pre-stimulation at pHo 5.0. The mutational analysis results, illustrated with current traces (Figs. 4*A* & 5*AB*), and displayed as bar graphs (Figs. 4*B* & 5*CD*), are as follows. 1/ The greatest impact on their ability to inhibit GLIC occurred with the removal of the pivot residue Arg77. The data on R77A was: in the presence of ACET 71.4 (± 16.6 %, n=4, P<0.001) % of control, with BUTYR 63.9 (±12.9, n=5, P<0.001) %, and with CROTON 65.6 (± 6.3, n=5, P<0.001) %, *versus* 12.2 (± 5.3, n=15), 10.3 (± 5.1, n=11), and 11.9 (± 6.6, n=11) % of control respectively on WT GLIC. 2/ The intra-SU CBX-binding pocket residue Glu104 was next, as ACET NAM effect was strongly reduced on E104Q, with, in the presence of ACET 47.6 (± 18.9, n=4, P<0.001) % of the control current. E104Q had a significant impact on BUTYR (35.6 ± 8.2 % of control, n=4, P<0.001) and CROTON (26.1 ± 5.3 %, n=4, P=0.002) NAM effects. 3/ The other intra-SU pocket mutation, Y102A, had little or no impact on ACET (16.1 ± 4.2 %, n=4, P=0.19), BUTYR (23.1 ± 10.1 %, n=4, P=0.006) and CROTON (14.2 ± 5.0 %, n=5, P=0.49) effects. 4/ Unexpectedly, the inter-SU CBX-binding pocket mutation R105A had little or no impact on the mono-CBX NAM effects, with the following data: with ACET 17.1 (± 4.9, n=6, P=0.067) % of control, with BUTYR 21.0 (± 5.6, n=5, P=0.002) %, and with CROTON 14.7 (± 5.0, n=5, P=0.42) % of control.

5/ The E181A substitution, known to have on GLIC no impact on CAFFE NAM effect (Prevost *et al*., 2013, pHo 5.5; Van Renterghem *et al*., 2023, pHo 5.0), and no impact on CROTON NAM effect (Alqazzaz *et al*., 2016, pHo 5.5), had no impact either on the NAM effect of ACET (Fig. 4), BUTYR and CROTON (Fig. 5) in our WT GLIC data at pHo 5.0. This data (no-impact on NAM effects) is related to the fact that, with the 4-carbon di-CBX, E181A not only abolished the PAM effect, but also revealed a unique di-CBX NAM effect (see Fig. 7 in Van Renterghem *et al*. (2023). 6/ Regarding the orthotopic site mutations, R133A had a minor significant impact in this protocol (see Discussion): with ACET, 22.7 (± 9.0, n=6, P=0.003) %; with BUTYR, 25.1 (± 5.1, n=4, P<0.001) %; with CROTON, 21.4 (± 6.6, n=4, P=0.029) % of control, as compared to the respective WT values,12.2 (± 5.3, n=15), 10.3 (± 5.1, n=11), and 11.9 (± 6.6, n=11) % of control already given above. E177A had almost no impact: with ACET, 14.9 (± 3.9, n=6, P=0.26) %; with BUTYR, 18.6 (± 7.2, n=5, P=0.019) %; with CROTON, 16.8 (± 4.8, n=5, P=0.16) % of control.

A “loose” pattern of mutational impact is therefore observed for the mono-CBX NAM effects, characterized by a weak impact (except for R77A) of every single mutation in the inter-SU pocket or its orthotopic entrance, or in the intra-SU (vestibular) CBX-binding pocket.

### Part 2 – A double mutation in Loop **Ω** favours positive modulation

#### 2-1 / The D86A-D88A pre-β5 double mutation has a week loss of function impact on proton-elicited activation

The pre-β5 strand, lining the extracellular vestibule lumen, is also part of the wall of the intra-SU CBX binding pocket on the axial side of GLIC ECD [opposite to Arg77 and the inter-SU CBX-binding pocket] (Sauguet *et al*., 2013, Fourati *et al*., 2015, 2020; see Fig. 6*A*). Indeed, the pre-β5 strand includes Arg85, whose side-chain belongs to the intra-SU pocket; Arg85 side-chain coordinates the intra-SU bound CBX if any, and is otherwise hold in place by Glu104, and a chloride (Fourati *et al*., 2020). Adjacent within the pre-β5 strand is the pair of aspartate residues, Asp86 and Asp88, pointing opposite, toward the vestibule lumen, where they bind cations in GLIC crystal structures (Sauguet *et al*., 2013, Fourati *et al*., 2015). The pre-β5 strand appears located somehow in-between the CBX-binding pockets and the gating machinery, as the (pre-)β5 – β6 “sandwich” ends down at the ECD – TMD interface with the β6 – β7 Loop, or Pro-Loop (Jaiteh *et al*., 2016), essential to gating. [In Eukaryote pLGICs, the Pro-Loop, stabilized by a disulfide bridge, is usually called Cys-Loop]. The pre-β5 strand in GLIC corresponds to the pLGICs Loop Ω defined by Hu *et al*., 2018, and analysed systematically in Eukaryote pLGICs structures by Brams *et al*., 2020.

Regarding GLIC activation by protons (characterized using the *Xenopus* oocyte expression system), the pre-β5 strand double mutation D86A-D88A (AA) produced a slightly loss of function (LoF) GLIC variant (Fig. 6*B*), with ΔpHo_50_ = -0.43 (± 0.19, n=3 [2 inj]), [in comparison to data recorded from oocytes expressing WT GLIC on the same day or the day before]. This data is consistent with the data published by Nemecz *et al*. (2017). To ensure that the impact of D86A-D88A on GLIC modulation by the mono-CBX was not a simple consequence of this LoF property, we included here another LoF mutant, N152A (ΔpHo_50_ = -0.31 ± 0.16, n=5 [2 inj]; Fig. 6*B*). Asn152 has a special interest, as it belongs to the inter-SU pocket, but has no contact with the inter-SU bound CBX when it is a mono-CBX molecule: Asn152 coordinates the second carboxyl group of the di-CBX molecule (Fourati *et al*., 2020). Consistently, in addition to the mono-CBXs, we tested here the di-CBXs and CAFFE.

**Figure 6A.**
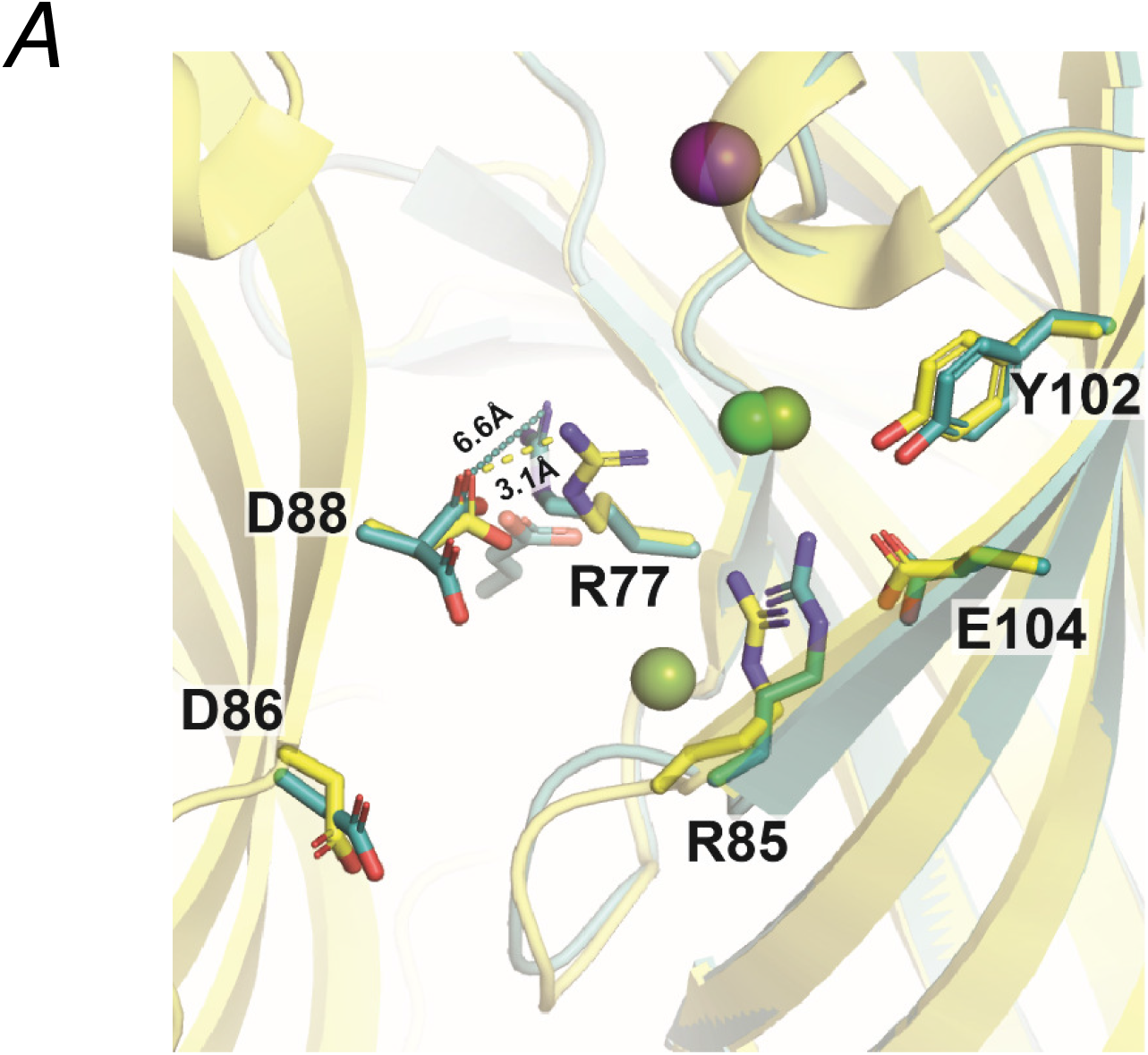
– The pre-β5 strand double mutation D86A-D88A (AA): *A*/ Location in GLIC structure. *A*, Location of Asp86 and Asp88 in GLIC crystal structure. Superimposed views from the Apo-GLIC (crystals grown in Phosphate buffer) structure (4qh5; *Yellow*), and the GLIC-CROTON co-crystal structure (6hji; *Dark cyan*). PDB references from Fourati *et al*. 2015 and 2020 respectively. Views are in GLIC ECD, from the axis of the pentamer towards periphery, with pre-β5 strand (n) on the left (bearing Asp86 and Asp88), and intra-SU pocket (n+1, *clockwise*) on the right. A chloride ion occupies the <u>intra-SU pocket</u> (“CBX-empty”) in the apo-GLIC structure (*Green sphere*) and the CROTON structure (*Yellow green sphere*). A second chloride (left to Arg85) is present in the CROTON structure (*Yellow green*). Whereas, in 6hji only, the <u>inter-SU pocket</u> (not represented) is occupied by a CROTON molecule, represented behind the Asp88 carboxyl group. A sodium ion is represented (*Light purple sphere* in Apo-structure, *Dark purple sphere* in CROTON-structure). An Asp88(n)-Arg77(n+1) ion bridge (3.1 Å) present in the Apo-structure is released in the CROTON structure (6.6 Å).

**Figure 6B.**
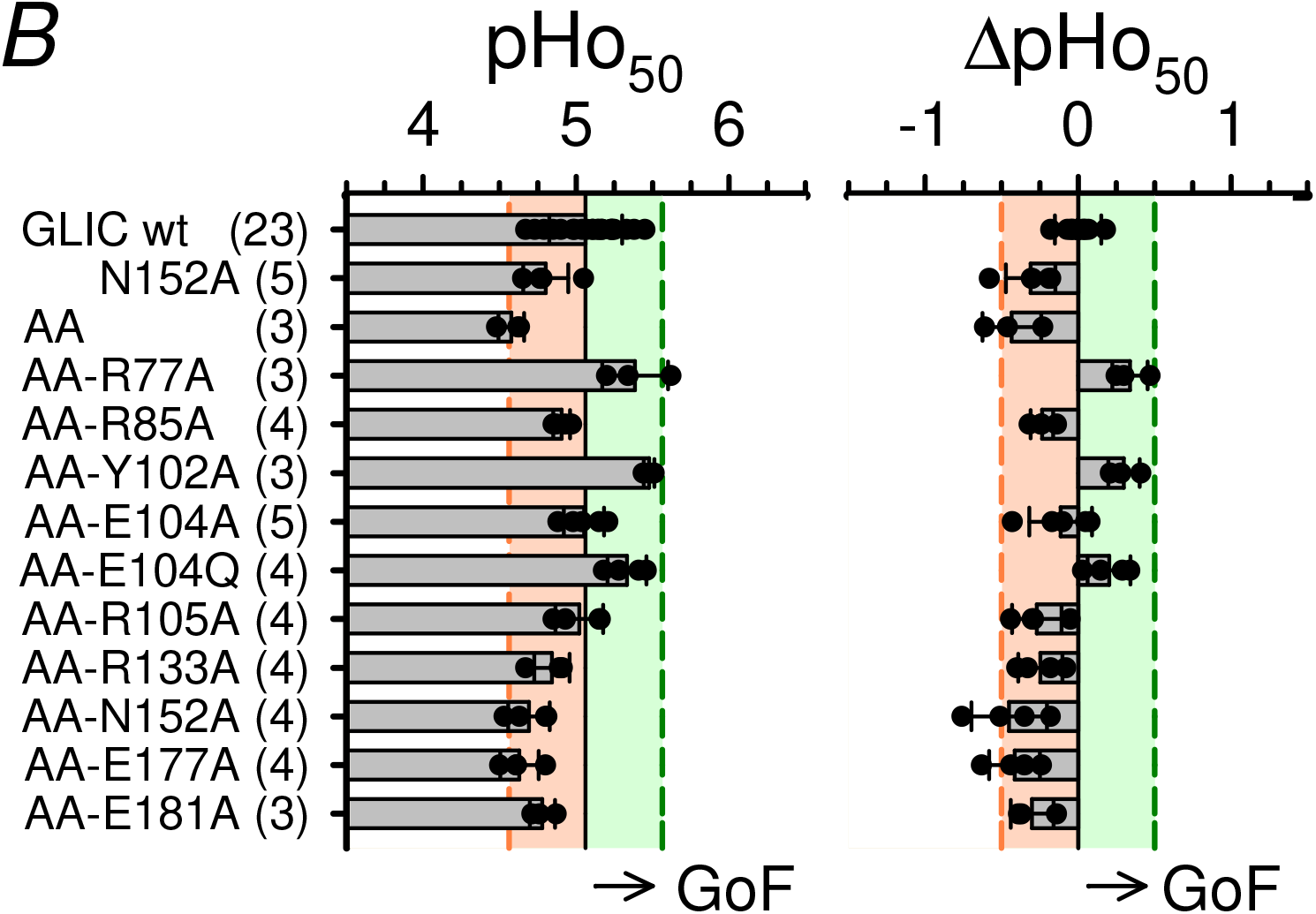
– The pre-β5 strand double mutation D86A-D88A (AA): *B*/ Low impact on GLIC proton sensitivity. *B*, Two-electrode voltage-clamp data obtained using the *Xenopus* oocyte expression system. Bar graphs showing values of the pHo corresponding to half maximal activation by low-pH extracellular solutions (pHo_50_, *Left*), and the difference between pHo_50_ values of mutant and WT GLIC (ΔpHo_50_, *Right*) (WT reference: recording on the same day or the day before, from 1 or 2 oocytes of the same batch, and in the same solutions, as the mutant considered). For each mutant is shown the data from 3 to 5 oocytes, coming from at least two injections. Pink and green coloured areas indicate the mutant inclusion criteria: ± 0.5 pH unit from the WT pHo_50_ value (EC_50_ *ratio* 0.3 to 3). In this representation, the data for a gain of function (GoF) variant goes right to WT.

#### 2-2 / The D86A-D88A pre-β5 double mutation favours compound-elicited positive modulation

Compound modulation of the D86A-D88A pre-β5 variant (AA), N152A mutant and WT GLIC, was investigated in the protocol with pre-stimulation at pHo 5.0 (Fig. 7). Surprisingly, CROTON effect (1 mM) was inverted on the pre-β5 double mutant, from an almost full inhibition at 13.6 % of control (± 6.1 %, n = 18) on the WT, into a strong potentiation at 232 % of control (± 59 %, n = 8, P <0.001) on the D86A-D88A mutant (Fig. 7*AB*). Such an inversion did not occur with BUTYR, the corresponding but saturated 4-carbon mono-CBX compound (Fig. 7*AB*). ACET and BUTYR remained inhibitors on AA, with no significant change for ACET (on wt: 12.2 ± 5.3, n = 15; on AA: 14.6 ± 3.0, n = 2, P = 0.54) and BUTYR, (on wt: 13.6 ± 5.9, n = 16 %; on AA: 18.0 ± 6.0, n = 5, P = 0.16).

**Figure 7.**
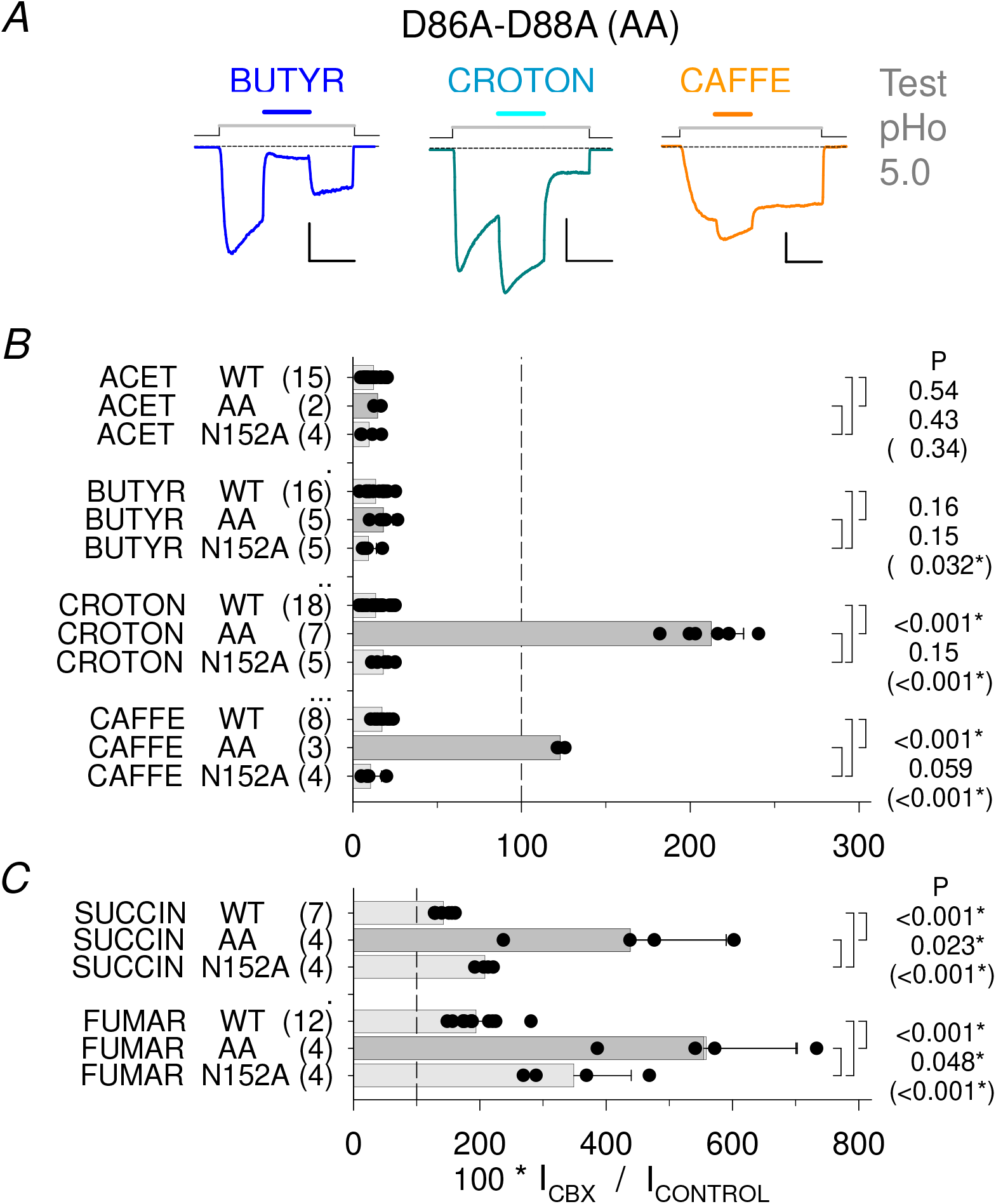
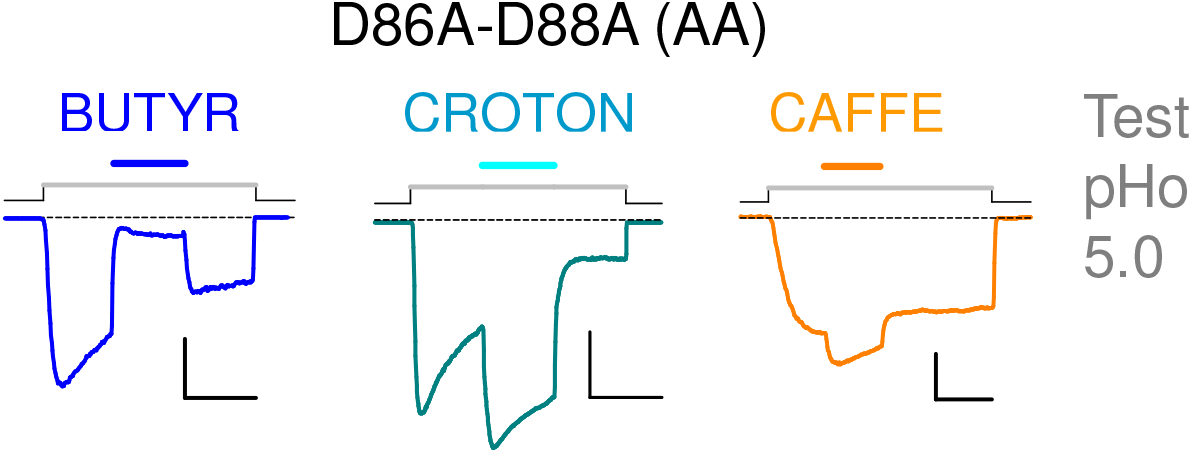
– The pre-β5 strand double mutation D86A-D88A (AA): major impact on compound-elicited modulation, and comparison with N152A. *A*, Current traces from three cells driven to express GLIC D86A-D88A (GLIC AA), showing the unchanged NAM effect of BUTYR (1 mM), the inverted, strong PAM effect of CROTON (1 mM), and inverted weak PAM effect of caffeate (0.1 mM), all at pHo 5.0. See Fig. 2CD or Fig. 5AB for the NAM effects of BUTYR and CROTON on WT GLIC at pHo 5.0. Scale bars, Current: 0.4 nA; Time: 20 s. *BC*, (*see* Figure 7BC *on previous page*). Bar graphs showing the current values (in % of control at pHo 5.0) obtained from cells expressing WT GLIC, or D86A-D88A, or N152A, in the presence of (*B*) a mono-CBX (ACET, BUTYR, CROTON; 1 mM), or CAFFE (0.1 mM), or (*C*) a di-CBX compound (SUCCIN, FUMAR; 10 mM). Statistics: Students’ T-test P values are indicated (top-down) for AA to WT, N152A to WT, and N152A to AA (in brackets) sample comparisons. <u>Comments</u>. On the pre-β5 variant AA, effects of CROTON and CAFFE are inverted, from near-full negative modulation on the WT, to positive modulation on AA, whereas BUTYR and ACET NAM effects are unchanged. SUCCIN and FUMAR PAM effects are strongly increased on the pre-β5 double mutant AA. N152A is a slightly loss-of-function variant, as is GLIC AA. However, in contrast with the impact of the pre-β5 double mutation AA, the N152A mutation has no impact on mono-CBX and CAFFE NAM effects (*B*),. Unexpectedly, N152A increases the di-CBX PAM effect (but less than AA) (*C*). Among variants with a slightly loss-of-function property regarding GLIC activation by protons (as shows AA), N152A was chosen because, in the GLIC-CBX co-crystal structures (Fourati *et al*. 2020), Asn152 side-chain is in contact with the second carboxyl group of the di-CBX compounds, but has no contact with the mono-CBX compounds.

CAFFE effect, however, was also inverted, from an almost full inhibition at 17.4 % of control (± 4.6 %, n = 8) on the WT into a mild potentiation at 123 (± 3 %, n = 3, P <0.001) % of control on AA (Fig. 7*B*). The other LoF mutation, N152A, had no significant impact on the effects of CROTON and the three other NAMs: for ACET 9.7 (± 5.8, n = 4, P = 0.43) % of control, for BUTYR 9.3 (± 4.8, n = 5, P = 0.15) %, for CROTON 18.1 (± 5.4, n = 5, P = 0.15) %, and for CAFFE 10.6 (± 6.5, n = 4, P = 0.059) % (Fig. 7*B*). The inverted effect of CROTON and CAFFE is therefore not determined by the weak LoF property, but specific to the pre-β5 double mutation.

The PAM effects of SUCCIN and FUMAR were strongly increased in the pre-β5 AA variant (Fig. 7*C*): for SUCCIN, 438 % of control (± 151 %, n = 4, P <0.001) on AA, *versus* 142 % (± 14 %, n = 7) on the WT; for FUMAR, 558 % of control (± 142 %, n = 4, P <0.001) on AA, *versus* 194 (± 36, n = 12) % on the WT (Fig. 7*C*). Unexpectedly, SUCCIN and FUMAR PAM effects were also *increased* on the N152A mutant (but to a lesser extent than in AA): on N152A, with SUCCIN, 208 % (± 12 %, n = 4, P_152vswt_ <0.001, P_152vsAA_ = 0.023); with FUMAR, 349 % (± 91 %, n = 4, P_152vswt_ <0.001, P_152vsAA_ = 0.048) (Fig. 7*C*).

#### 2-3 /Functional relevance of GLIC CBX binding pockets for CROTON PAM effect on the pre-**β**5 GLIC variant D86A-D88A: mutational analysis on an AA basis (pHo 5.0)

The residue dependency analysis (Figs. 4, 5) showed that single mutations in the CBX-binding pockets have a weak impact (except for R77A) [“loose” pattern] on the mono-CBX NAM effects, contrasting with their “all-or-none” impact on the di-CBX PAM effects (see Fig. 7 in Van Renterghem *et al*., 2023). As CROTON is converted into a PAM on the AA pre-β5 variant (Fig. 7*AB*), we decided to check the impact of adding single mutations in the CBX-binding pockets (on an AA basis). Would CROTON PAM effect on AA display a mono-CBX/NAM type “loose” pattern (according to compound structure)? Or would it display a di-CBX/PAM type “all-or-none” pattern of impact (according to compound effect)? We therefore added the pre-β5 double mutation to every CBX-binding pocket single mutant.

The triple mutants were first characterized for their sensitivity to pHo (Fig. 6*B*). Expressed in *Xenopus* oocytes, they were all functional, and presented pHo_50_ values within the mutant inclusion *criterium*, *i*.*e*. within ±0.5 pH unit from the pHo_50_ value measured for WT GLIC [with the same batch of oocytes and solutions]. Therefore, the triple mutants were all included in the study.

CROTON was tested in whole-cell patch-clamp, in the conditions previously used for the CBX-pocket single mutants. CROTON effect on the [AA + CBX-pocket] triple mutants was compared to its effect on the AA double mutant (Fig. 8). Adding anyone of the binding pocket mutations fully abolished CROTON PAM effect, despite the presence of the pre-β5 double mutation (Fig. 8*AB*). Therefore, CROTON PAM effect (inverted on AA, *vs* WT) clearly displays a di-CBX/PAM type pattern, with an “all-or-none” impact of every binding pocket mutation [according to a CBX PAM effect]. Here however, with AA, CROTON ability to positively modulate GLIC being fully abolished (by any binding pocket mutation), its ability to negatively modulate GLIC is then revealed. And the recovered negative modulation by CROTON displays the mono-CBX/NAM type “loose” pattern of impact [as now compared with WT GLIC] (Fig. 8*B*), similar to the pattern observed when adding the single mutations to GLIC WT (see Figs. 4,5): Each mutation had a weak impact; greatest impact observed with R77A (removing the pivot); next greatest impact with an intra-SU pocket mutation; weak significant impact of orthotopic site mutations R133A or E177A, and inter-SU pocket mutation R105A; no impact of E181A [*i*. *e*. according to a mono-CBX structure].

Finally, the N152A mutation, which had no impact on CROTON NAM effect (WT basis; Fig. 7*B*), but increased SUCCIN and FUMAR PAM effects (WT basis; Fig. 7*C*), also increased CROTON PAM effect (AA basis; Fig. 8*C*). The potentiating impact of N152A on an AA basis was significant (381 ± 189 %, n = 13, P_AA152vsAA_ = 0.0037, *vs* 183 ± 28 %, n = 10 for AA, as mentioned). However, as the values for CROTON effect on AA-N152A were noticeably dispersed, we decided to plot the PAM ratio in % (= 100*I_CROTON_/I_CONTROL_) according to the control current amplitude (absolute value). This representation shows that the PAM ratio (%) decreases with increasing control amplitude. It also makes clear that the value is consistently increased in AA-N152A *vs* AA.

## Discussion

### Short chain mono-CBX as negative modulators of allosteric transitions in GLIC

We report that short-chain, saturated mono-CBX compounds are negative modulators of the allosteric transitions in GLIC. The NAM effect of ACET, PROPION, BUTYR, or VALER (2-to 5-carbon compounds) occurs with no influence of the carbon chain length on the concentration – effect curve (Fig. 2). CROTON was previously published to negatively modulate GLIC (Alqazzaz *et al*., 2016). Our data shows that the double bond present in CROTON, not in BUTYR, has no influence on the concentration – NAM effect curve (Fig. 2*CD*). These properties contrast with the fact that a 4-carbon di-CBX is a better PAM than a 3- or 5- carbon di-CBX, and FUMAR (with double bond) a better PAM than SUCCIN (see Figs. 1*AB* & 2*A*-*D* in Van Renterghem *et al*., 2023). And both data are consistent with the previously published GLIC-CBX co-crystal structures (Sauguet *et al*., 2013, Fourati *et al*., 2015, 2020), showing that the mono-CBXs are bound by a single end of the molecule (the carboxyl group), whereas the di-CBX compounds are bound by both ends.

Mutations located away from the pore or vestibule lumen (R77A, E104Q) abolish the major part of the mono-CBX inhibitory effect (Figs. 4, 5), suggesting that inhibition is not due to a channel block or another permeation mechanism. The fact that SUCCIN PAM effect overcomes ACET inhibitory effect (Fig. 1*B*) also excludes a permeation mechanism. In addition, the D86A-D88A double mutation inverts inhibition to a potentiation (CROTON and CAFFE, Fig. 7). The data therefore demonstrates that, at least for CROTON and CAFFE, we are actually dealing with a compound-binding elicited negative modulation of GLIC gating.

### Inhibition by CBX compounds and GLIC current decay

Given their low p*K*a values, carboxylic acids have been reasonably chosen as low-pH buffers in some functional studies of GLIC. The mono-CBX NAM property then produces a (relatively) fast GLIC current decay (due to compound inhibition, as in Fig. 1*A*), which may be misinterpreted as low pHo related GLIC desensitization. CBX compounds cannot be used as pH-buffers in functional studies of GLIC, and previously published reports using ACET as buffer need to be reinterpreted. The tri-CBX citrate, used by some authors, had no effect on GLIC current, but produced some instability in the recording. We chose to keep-on with Good’s buffers (thought to be non-membrane-permeant), despite the fact that the lowest p*K*a value available (near 6.2 for MES) gives a poor buffering capacity at the low pHo values used with GLIC. This point is counter-balanced by the use of continuous extracellular solution flow in electrophysiological recordings.

Regarding GLIC desensitization, our data shows that, in the absence of a NAM compound, low pHo induced GLIC current decay is very slow (Fig. 1*A* Currents in *Grey*). As mentioned by other authors, it is also very variable. Van Renterghem *et al*. (2023, Fig. 3 and text) proposed that a progressive (and variable), low pHo-induced, *drop in intracellular pH* may be the major determinant of GLIC current decay kinetics (at least in the absence of compound). According to the preliminary data published by Hilf & Dutzler (2009), with pHi = 4, GLIC is no more activated by its “agonist”, a pHo 4 extracellular solution (the point, however, requires further investigation). Would GLIC at low pHi correspond to its desensitized state? Whether a desensitized state exists or not in GLIC becomes a question of definition. And whether a desensitized state (defined as in Eukaryote pLGICs) may be favoured by orthotopic binding of a PAM compound, such as FUMAR, remains an unanswered question.

### A double bond favours positive modulation of allosteric transitions, and is associated with exclusive inter-SU binding in the structures

Regarding the 4-carbon compounds, the double bond favours the PAM effect of a di-CBX (FUMAR *vs* SUCCIN; see Figs. 1*AB* & 2*CD* in Van Renterghem *et al*., 2023), not the NAM effect of a mono-CBX (BUTYR and CROTON have indistinguishable NAM effects on GLIC WT; Figs. 2*CD*, 5, 7*B*). GLIC binding sites, however, are able to distinguish CROTON from BUTYR (at least in the pre-β5 variant), since CROTON modulatory effect is then inverted, whereas BUTYR NAM effect is unchanged (Fig. 7*AB*). Here, a double-bond is required for the 4-carbon mono-CBX to be converted into a PAM on the pre-β5 variant. As is the case for CROTON and FUMAR, the CAFFE molecule has a *trans* double bond in *alpha* to its carboxyl group. Consistent with our conclusion, CAFFE effect was also inverted to a PAM on the pre-β5 variant (Fig. 7*AB*). From these functional data, we conclude that a double bond has no impact on NAM effects, but favours PAM effects.

In the crystal structures of GLIC-CBX complexes (Sauguet *et al*., 2013, Fourati *et al*., 2020; see Table 1), the 2- or 3-carbon mono-CBX compounds (ACET, PROPION) and the saturated 4-carbon di-CBX (SUCCIN) are found occupying the two CBX-binding pockets. In contrast, the 4-carbon compounds with a double bond (CROTON, FUMAR), both occupying the inter-SU pocket, are absent from the intra-SU pocket (see Table 1 & Fig. 6*A*), suggesting that the vestibular pocket cannot handle the compounds with a double bond. No co-crystal structure is available for BUTYR or CAFFE; but from the available structures, it may be expected that the saturated 4-carbon mono-CBX would probably be found in the two pockets (as ACET, PROPION and SUCCIN), but CAFFE (with its C2-C3 double bond) only in the inter-SU pocket.

### The “all-or-none” pattern of mutational impact for CROTON inverted effect supports the model with inter-SU binding and vestibular control

The spectacular inversion of CROTON effect (not BUTYR effect) on the D86A-D88A pre-β5 variant (Fig. 7*AB*), together with the exclusive inter-SU binding of CROTON in the available structure (see Fig. 6*A*), support the involvement of an (exclusive) inter-SU binding of CROTON in this positive modulation of the allosteric transitions by a mono-CBX. Our residue-dependency analysis shows that the CROTON PAM effect (on the AA basis, Fig. 8*AB*) is labile, as much as SUCCIN or FUMAR PAM effects on WT GLIC (see Fig. 7 in Van Renterghem *et al*., 2023), since CROTON PAM effect was fully abolished by any single mutation either in the inter-SU pocket, or in the intra-SU pocket as well. This “all-or-none” pattern is consistent with the view that positive modulation by CROTON requires inter-SU binding, and integrity of the vestibular region corresponding to the intra-SU pocket. The model proposed for the di-CBX PAM effect and CAFFE NAM effect on wt GLIC (see Fig. 9 in Van Renterghem *et al*., 2023) now applies also to CROTON PAM effect on the pre-β5 variant: binding to the inter-SU site, and involvement of the vestibular region.

It is noticeable that the E181A mutation, when applied on the GLIC AA basis, suppresses CROTON PAM effect (Fig. 8*AB*), whereas, applied on a WT GLIC basis, E181A does not suppress CROTON NAM effect (Fig. 5*B*, *D*): a mutation in the orthotopic site Loop C suppresses the PAM effect, not the NAM effect, of the same compound. Given that CROTON is thought to bind exclusively to the orthotopic/inter-SU site, the data leads to the conclusion that Glu181 is specifically required for (compound-elicited) positive, not negative modulation.

### Mechanism for the mono-CBX NAM effect

The situation is different and more difficult to understand with the mono-CBX NAM effect. Indeed, the residue-dependency in the CBX binding pockets is very similar for ACET, BUTYR and CROTON NAM effects on WT GLIC (Figs. 4, 5). In all these cases, the “loose” pattern of mutational impact is completely different from the “all-or-none” pattern discussed above. With the mono-CBX NAM effects (Figs. 4, 5), apart for R77A, every mutation has only a weak significant impact, or no impact at all. The greatest impact of removing Arg77 suggests that some coupling between inter- and intra-SU pockets is involved here too, suggesting a necessary inter-SU binding. However, single mutations in the inter-SU pocket (R105A) or its orthotopic site entrance (R133A, E177A) have no impact at all (as E181A) or a weak significant impact. Significant mutational impact is observed in the intra-SU pocket (Tyr102, Glu104), but it is weak too. Here, the model with inter-SU binding and vestibular control is under questioning.

We therefore propose the following interpretation (Fig. 9, also supported by crystallographic data): the model with the inter-SU pocket as main binding site still applies, with required integrity of the vestibular pocket. But negative modulation is “easy to reach” and occurs even when binding contacts or residue interactions are poor, whereas positive modulation requires stringent conditions, so that positive modulation is fully lost when a single residue is missing in one of the pockets. This interpretation is supported by the fact that, when CROTON PAM effect (AA basis) is abolished by any single mutation in the CBX-binding pockets (“all-or-none” pattern), then CROTON does not become inactive (as were the di-CBX on most single-mutants (WT-basis; see Fig. 7 in Van Renterghem *et al*., 2023). But CROTON recovers its NAM property (Fig. 8*AB*), with the NAM-specific “loose” pattern of residue dependency.

**Figure 8AB.**
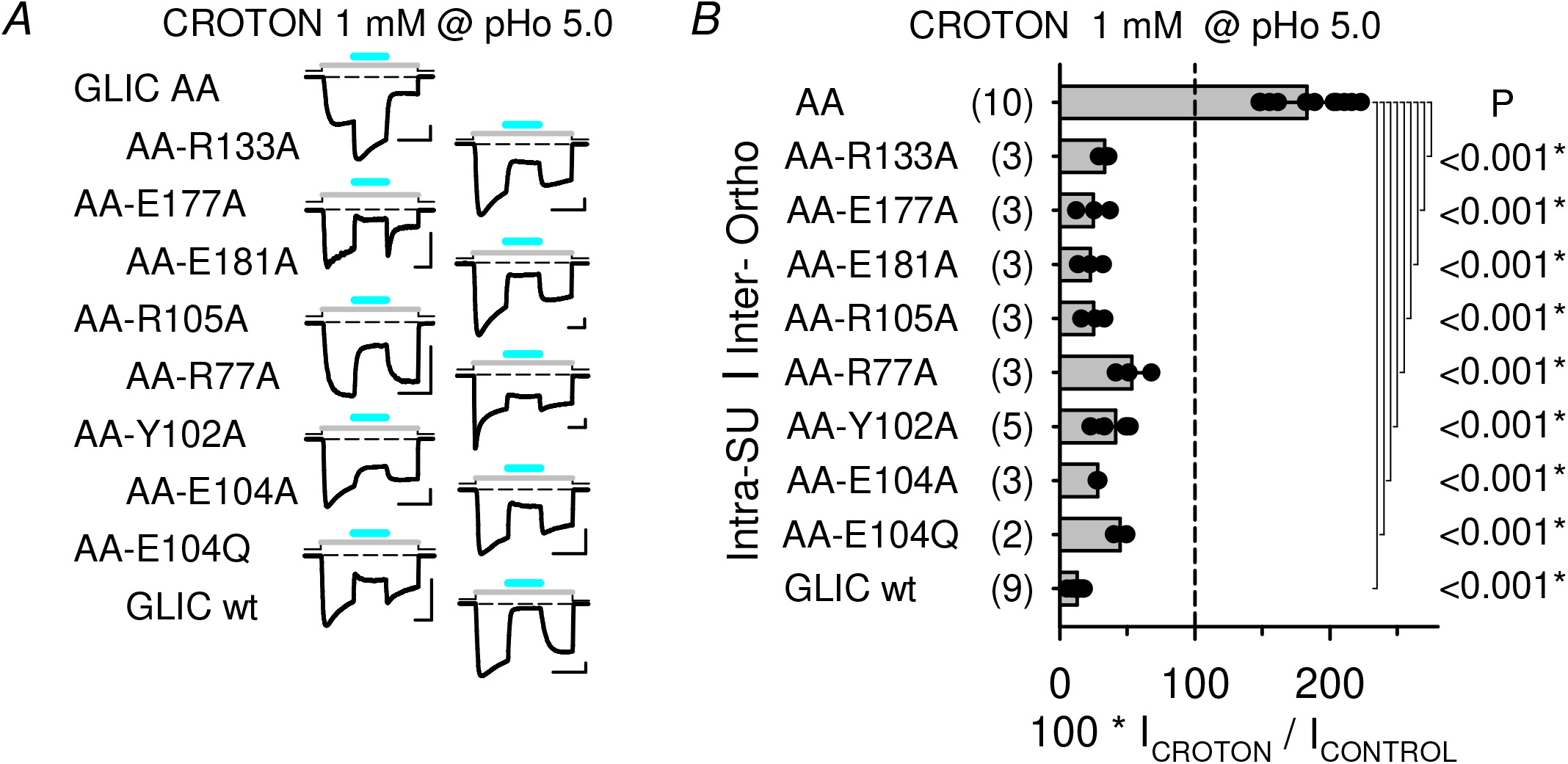
– CBX-pockets single mutations added to the AA variant: impact on CROTON inverted, PAM effect. *A*, Representative current traces illustrating CROTON tests (1 mM at pHo 5.0) on GLIC AA, triple mutation GLIC variants, and WT GLIC, as indicated. Scale bars, Current: 0.2 nA (AA-E177A), 1 nA (R77A), or 0.4 nA (others); Time: 10 s. *B*, Bar graph of current in the presence of CROTON (1 mM; in % of control at pHo 5.0), on the pre-β5 double mutant D86A-D88A (AA), on the triple mutation GLIC variants, and on WT GLIC. Statistics: each triple mutant sample of data was compared to the AA double mutant sample. <u>Comments</u>: Regarding the CBX binding pocket single mutations, CROTON PAM effect on the pre-β5 double mutant shows the “all-or-none” pattern of residue dependency, previously observed with the di-CBX PAM effect (see Fig. 7 in Van Renterghem *et al*., 2023): any CROTON PAM effect is lost with anyone of the single CBX-binding sites mutations. However, with the CBX-pockets mutations on the AA basis, a CROTON NAM effect reappears, showing a “loose” pattern of residue-dependency, as with ACET, BUTYR and CROTON NAM effect on WT GLIC (see Figs. 4, 5).

**Figure 8CD.**
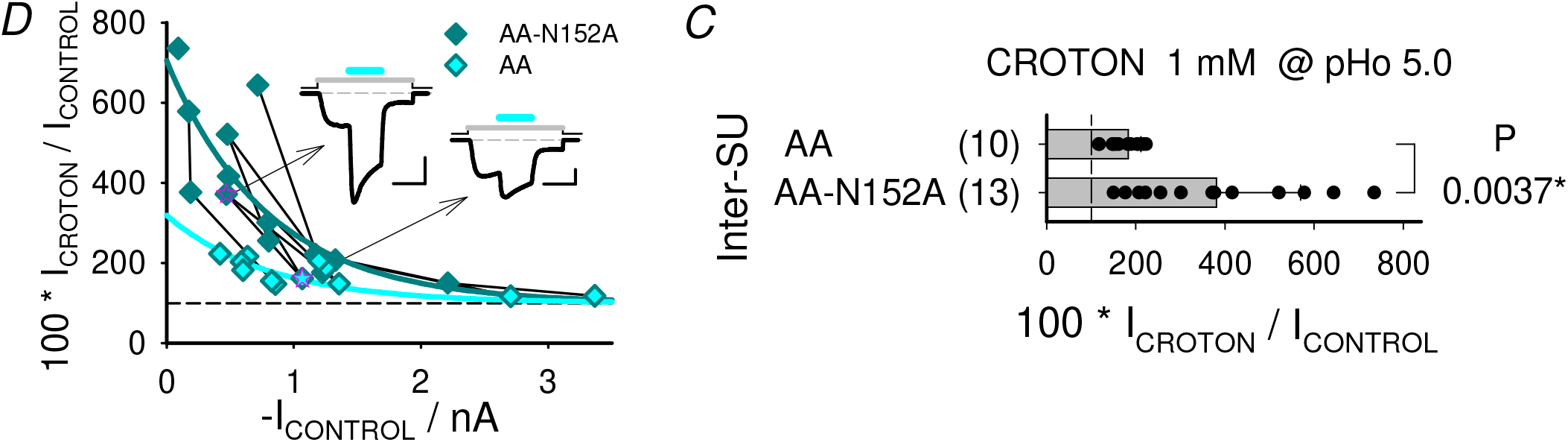
– CBX-pockets single mutations added to the AA variant: impact of N152A (added to AA) on CROTON inverted, PAM effect. *C*, Bar graph of current in the presence of CROTON (1 mM; in % of control at pHo 5.0), on the pre-β5 double mutant D86A-D88A without (AA) and with the N152A mutation (AA-N152A). Statistics: Student T-test compares AA-N152 and AA samples of data. *D*, The CROTON data shown in *C* was plotted against the control current amplitude (absolute value) for AA-N152A (*Dark Green* diamond), and AA (*Cyan* diamond). Data points obtained on the same day of recording are joined by a black line. The current traces displayed as *Inset* were recorded, on the same day, from an AA-N152A cell (*Left* trace) and an AA cell (corresponding data points spotted in pink in plot). Scale bars, Current: 0.4 nA; Time: 10 s. The PAM ratio (in %) decreases with increasing current amplitude. A single exponential decay function (*Dark Green* line) (decaying toward y0 = 100 %, fixed) was fitted to the AA-N152A data, giving a decay constant tau = 800 nA, and an amplitude (at 0 nA) of 604 %. The AA data, arbitrarily fitted (*Cyan* line) with the same decay constant value (800 nA, fixed) gave an amplitude (at 0 nA) of 219 %. <u>Comments</u>: The data in *C*, and its further analysis in *D*, clearly show that the N152A mutation increases CROTON PAM effect on the AA variant (as N152A unexpectedly increases the di-CBX PAM effects on wt GLIC, Fig. 7C).

**Figure 9.**
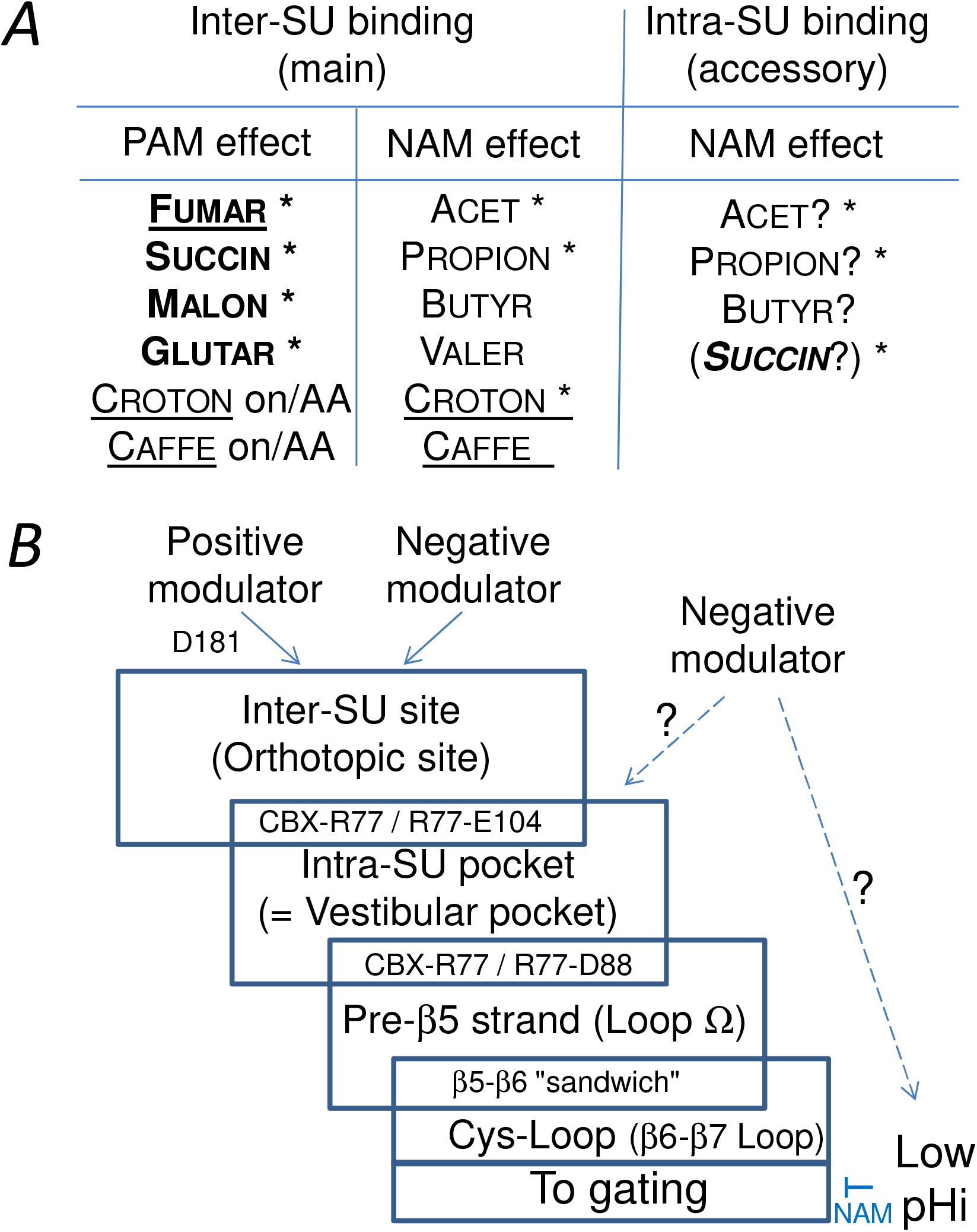
– A mechanistic hypothesis compatible with crystallographic and electrophysiological CBX data on GLIC. Co-Crystal structures: Sauguet *et al*. 2013, Fourati *et al*. 2015, 2020. Electrophysiology: Van Renterghem *et al*. (2023) and present Report. (*See details next page*).

An additional hypothesis may help to explain the mono-CBX “loose” impact data: intra-SU (accessory) binding of saturated mono-CBX compounds might be sufficient to produce a negative modulation, at least when the inter-SU pocket (or its orthotopic site entrance) is not intact. Indeed, in the available structures, ACET and PROPION are found in both inter-SU and intra-SU pockets. It is also the case for SUCCIN. But SUCCIN has no effect at all on the inter-SU mutants, suggesting that intra-SU binding alone cannot promote positive modulation. We therefore add to the “in series” model an accessory vestibular (downstream) binding site, putatively involved in negative modulation (Fig. 9). Although this point is *compatible* with our data, it is not *demonstrated* by our data.

We finally raise here the hypothesis (Fig. 9) that some intracellular acidification may occur following transmembrane permeation of the acid forms of extracellularly applied mono-CBXs (see Thomas, 1974): a drop in pHi would contribute to negatively modulate GLIC activity (see Fig. 3 in Van Renterghem *et al*., 2023). Although we have no data showing the occurrence of a drop in pHi, and although we record using a patch-pipette filled with a strongly proton-buffered solution (10 mM HEPES + 10 mM of the tetra-acido-basic Ca^2+^-buffer BAPTA; pH 7.3), we do not exclude some contribution of sub-membrane acidification to the mono-CBX NAM effect, which would end up in “loosening” the mutational impact pattern, bringing some “blindness effect” in the analysis of the receptor-mediated mechanism, in the case of mono-CBX NAM effects. Indeed the “slow” recovery from ACET inhibition (see current traces in Figs. 1*B*, 2*A*, 4*A*) is compatible with the “slow” recovery from pHi lowering. But the consistently “fast” recovery from BUTYR or CROTON inhibition (see traces in Figs. 2*C*, 5*AB*) excludes a major contribution of intracellular acidification to BUTYR and CROTON inhibition. And, finally, the mutational impact pattern for ACET (Fig. 4), which is not “more loose” than the pattern for BUTYR and CROTON NAM effects (Fig. 5), but very similar, suggests that the contribution of a pHi effect may be minor with ACET as well, comforting a receptor-mediated mechanism for all mono-CBX NAM effects.

**Figure 9A.**
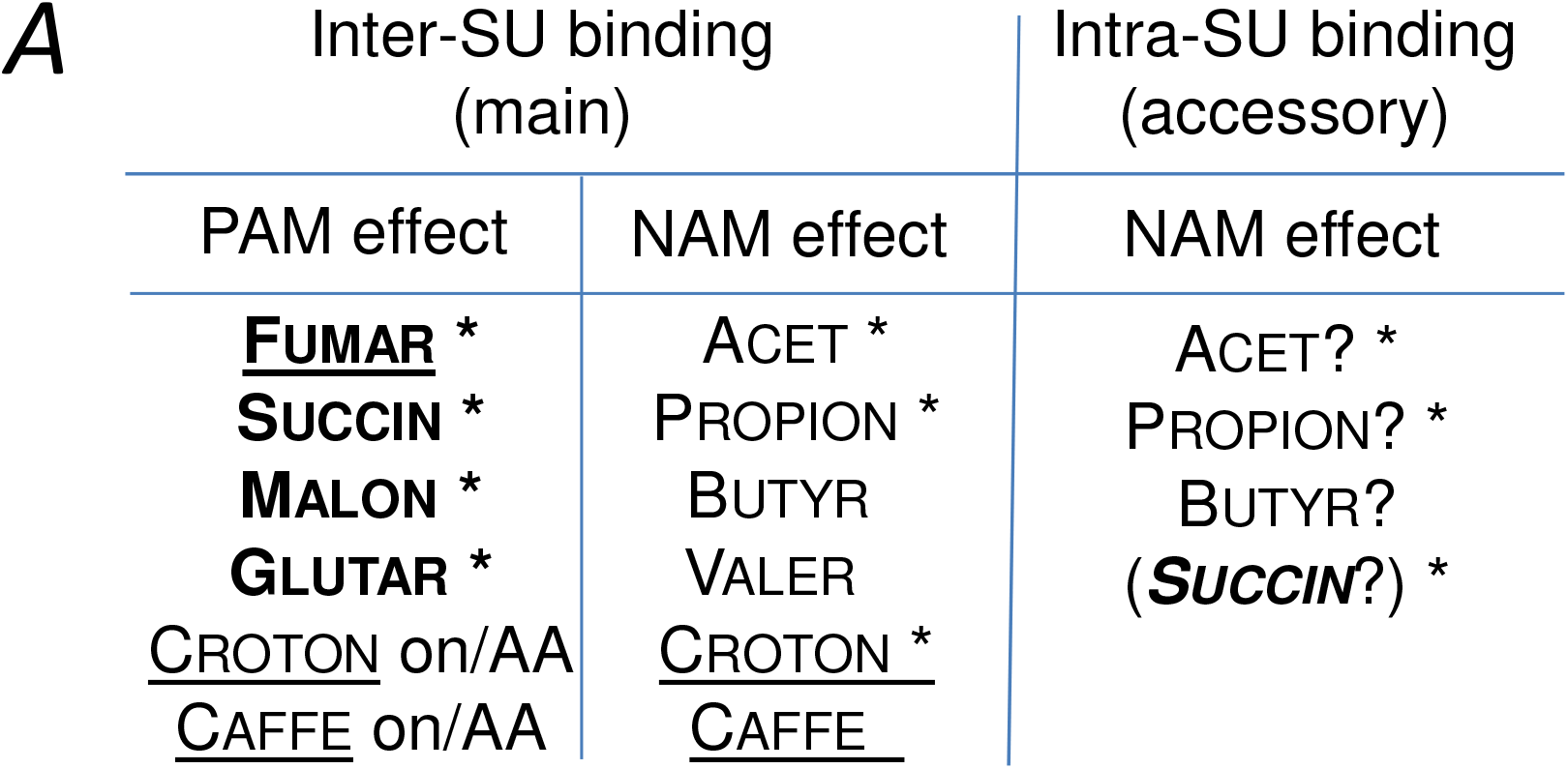
– A mechanistic hypothesis compatible with crystallographic and electrophysiological CBX data on GLIC. Co-Crystal structures: Sauguet *et al*. 2013, Fourati *et al*. 2015, 2020. Electrophysiology: Van Renterghem *et al*., (2023), and present Report. *A*, Codes: * *Star*, presence observed in a co-crystal structure; ?, suggested, not demonstrated, from functional data; *Bold*, a di-CBX; *Underlined*, with a *trans* double-bond (binds only to inter-SU site); on/AA, effect on the pre-β5 double mutant AA; *In brackets*, suggested secondary NAM influence of SUCCIN, limiting its PAM effect. *ERRATUM,* Figure 9A: We apologize for <u>an error in the previous BioRxiv version</u> of this report. Malonate and glutarate were represented by mistake in the intra-SU pocket. Malonate and glutarate are both present only in the inter-SU pocket (as shown here) in the co-crystal structures published by Fourati *et al*. 2020.

**Figure 9B.**
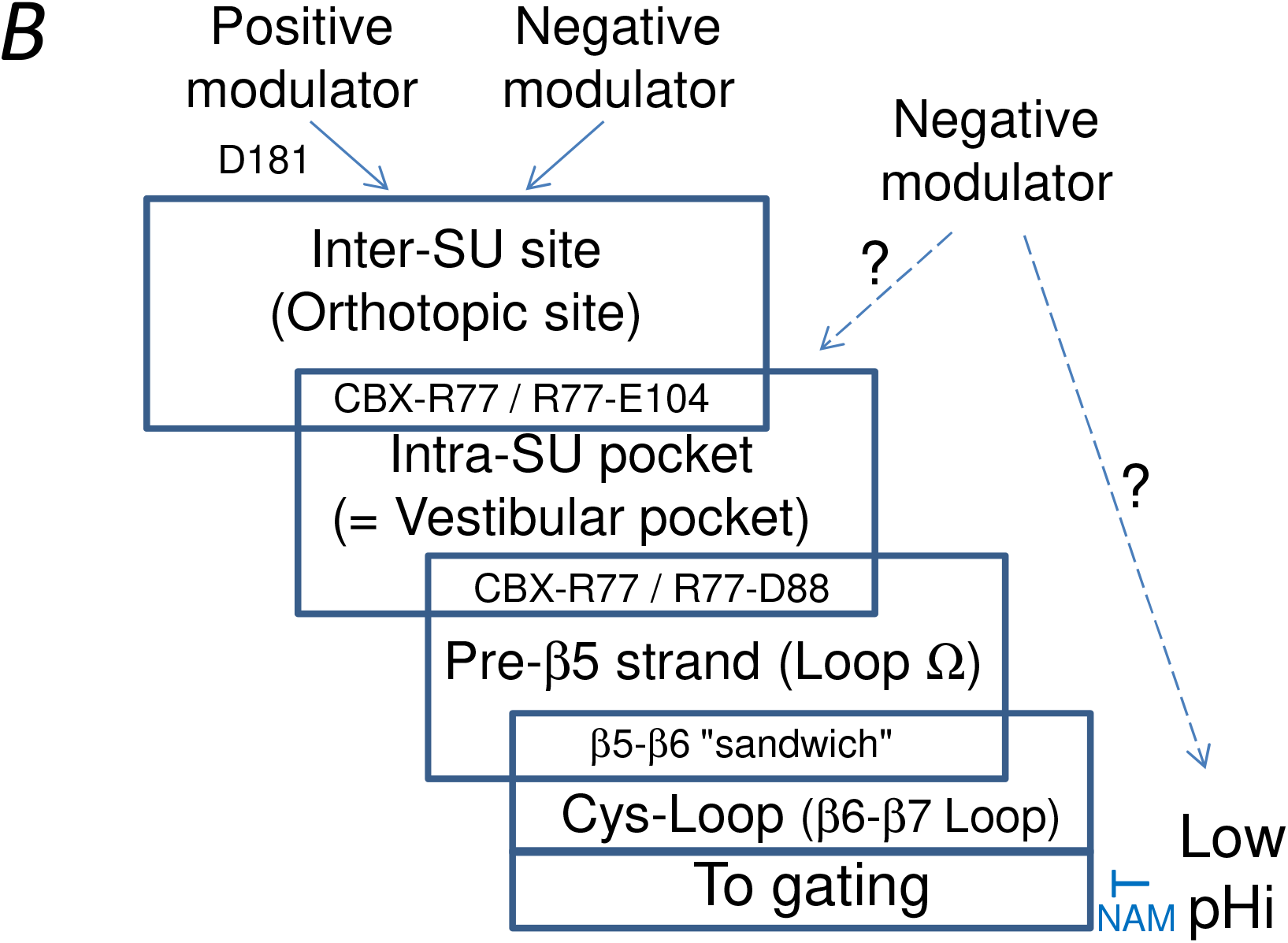
– A mechanistic hypothesis compatible with crystallographic and electrophysiological CBX data on GLIC. Co-Crystal structures: Sauguet *et al*. 2013, Fourati *et al*. 2015, 2020. Electrophysiology: Van Renterghem *et al*. (2023) and present Report. *B*, We suggest that only inter-SU binding may lead to a PAM effect. Caffeate NAM effect is thought to occur by binding to the inter-SU site. The mono-CBXs are thought to modulate GLIC by binding to the inter-SU site. But the data does not exclude a contribution of intra-SU binding to negative modulation. Low-intracellular pH (pHi) was shown to negatively modulate GLIC by Van Renterghem *et al*., (2023). We do not exclude the hypothesis that some mono-CBX-elicited drop in sub-membrane intracellular pH (pHi) may contribute to the negative modulation of GLIC in our recording conditions, thus “loosening” the mutational impact pattern for the mono-CBX NAM effects.

### Impact of R133A on CROTON NAM effect

Some discrepancy appears between our data and the data published by Alqazzaz *et al*. (2016), regarding R133A impact on CROTON effect (for WT GLIC). We found that the quantitative difference in the data has a kinetic explanation: with BUTYR or CROTON, inhibition is slowed down in R133A, as visible in our current traces using a protocol with pre-stimulation (Fig. 5*AB*). In our protocol with pre-stimulation, the current in the presence of CROTON is measured after 20 s of CROTON action, *i*.*e*. at the steady-state of inhibition (on both WT and R133A), leading to the conclusion that R133A has a weak impact on inhibition by CROTON. In the direct protocol, in WT GLIC, inhibition by BUTYR or CROTON occurs during activation by protons (as in Fig. 1*A*), whereas in R133A, delayed inhibition occurs as a decay following the peak of activation by protons, so that measuring the peak current leads to the conclusion that inhibition is almost abolished in R133A. We confirmed this explanation by performing patch-clamp recordings in the direct protocol, with BUTYR and CROTON: then (as Alqazzaz *et al*.) we found that peak current inhibition is almost fully abolished in R133A, with both BUTYR and CROTON. Both protocols are valid, they give slightly different informations.

In the crystal structures published in 2020 by Fourati *et al*., Arg133 (and Glu177) belong to the peripheral entrance of the deeper inter-subunit CBX-binding pocket. The authors note that “Arg133 partially obstructs the putative orthosteric pocket”. Here we propose that Arg133, located in the access corridor (and most probably positively charged) is actually *required* for a fast access of (negatively charged) BUTYR or CROTON to their more deeply buried (inter-SU) binding site. With this explanation, R133A strong impact in the direct protocol (Alqazzaz *et al*., 2020), and R133A weak impact in the protocol with pre-stimulation (this report, Fig. 5) are both consistent with the relatively long distance between Arg133 and the CROTON molecule in the crystal structure. The general consistency of the point commented here comforts again the idea that CROTON and BUTYR act by binding to the inter-SU site.

### The unexpected impact of N152A

In the GLIC-CBX co-crystal structures (Fourati *et al*., 2020), Asn152 coordinates the second carboxyl group of the di-CBX molecules bound in the inter-SU pocket, and does not contact mono-CBX molecules, [while, at the other end of the di-CBX molecule, carboxyl group one is in contact with Asp181, Arg105, and Arg77, as with the mono-CBXs]. We were then expecting that removing Asn152 would abolish di-CBX effects, and leave mono-CBX effects unchanged. Indeed, the N152A mutation has *some impact* on the di-CBX PAM effect (Fig. 7*C*), and *no impact* on the mono-CBX NAM effects (Fig. 7*B*), consistent with Asn152 *contacting* or *not contacting* the compound, respectively. But SUCCIN and FUMAR PAM effects were unexpectedly *increased* in N152A (Fig. 7*C*). This suggests that Ala substitution of Asn152, or binding a di-CBX on Asn152, may result in applying locally common forces involved in promoting positive modulation of the allosteric transitions. This hypothesis is supported by the fact that adding the N152A mutation to the pre-β5 variant also increased CROTON (inverted) PAM effect (Fig. 8*CD*), even though Asn152 does not contact the CROTON molecule in the available co-crystal structure. [It may be noted that the facilitating impact of adding N152A to the D86A-D88A variant is not due to the mild LoF property of the AA-N152A variant (regarding activation by protons), since the N152A, AA, and AA-N152A variants show approximately equal ΔpHo_50_ values (Fig. 6*B*)]. From our very unexpected data with N152A and AA-N152A, we conclude that Asn152, with its local interactions in the protein, exerts in GLIC ECD some resistance to compound-elicited activation. And that usually accepted interpretations, in a mutational structure-function study, are not always the right ones.

### Compound-elicited activation involves a motion of the pre-**β**5 strand

A related conclusion may be derived from our observation that Ala-substitution of the double ring of pre-β5 Asp residues strongly favours a PAM (*versus* NAM) effect of compound binding (Fig. 7): local interactions of Asp86 and/or Asp88 exert in GLIC ECD some resistance to compound-driven activation. In contrast, the D86A-D88A double mutation does not favour low pHo-controlled activation, as the AA variant is not a gain of function (GoF), but a mild LoF regarding activation by protons (Fig. 6*B*, and Nemecz *et al*., 2017).

Regarding the position in the agonist (proton) concentration – activation curve, our tests for CBX modulation in the WT or mutants are done at a constant pHo value of 5.0, or proton activity equal to 10^-5^, which is near proton EC_50_ on WT GLIC, and <EC_50_ on GLIC AA (slightly LoF). This may lead to some increased PAM effect on AA *vs* WT, as predicted by Rubin & Changeux (1966) in the Monod, Wyman & Changeux allosteric model (Monod *et al*., 1965). But the AA variant, and the N152A mutant, have equal ΔpHo_50_ values, whereas facilitation of the di-CBX PAM effect is larger with AA (Fig. 7*C*). Moreover, CROTON and CAFFE effects are inverted on AA, not on N152A (Fig. 7*B*), showing that something specific occurs with the pre-β5 variant (*vs* N152A), something regarding compound-driven activation.

### The Arg85(n)-Arg77(n), and Asp88(n)-Arg77(n+1) ion bridges: two Arg77 anchorings are released in CBX-bound structures

Comparison of the crystal structures for apo-GLIC (Fourati *et al*., 2015) on one side, and, on the other side, for GLIC-acetate (Sauguet *et al*., 2013) and other GLIC-CBX complexes (Fourati *et al*., 2020), leads to remarkable observations regarding Arg77 (at the border between inter- and intra-SU pockets). Arg77 interactions with Arg85 (pre-β5 residue pointing to the intra-SU pocket), and with the Asp86-Asp88 pair (pre-β5 residues pointing together to the vestibule lumen) are released in CBX-bound structures.

In an axial view of the pentamer, looking from extracellular to intracellular sides, we number subunits n to n+1 clockwise (which is, in a view from the vestibule lumen to the periphery of the pentamer, n to n+1 left to right). In the apo-GLIC structure (4qh5; Fourati *et al*., 2015), the side-chain of the pre-β5 residue Arg85(n) points to the periphery of the pentamer, and, within a subunit, interacts in the middle of the intra-SU pocket with the side-chain of Arg77(n) [in its apo-GLIC orientation]. The pre-β5 (n) next residues, Asp86 and Asp88, have their side-chains pointing towards the vestibule lumen, on the (n) to (n+1) side of the intra-SU pocket (n) (see Fig. 6A). And Asp88(n) side-chain is involved in an ion bridge with the side-chain of Arg77(n+1) [from the adjacent subunit, also in its apo-GLIC orientation] (Fig. 6A). Therefore, Arg77(n), in its apo-GLIC orientation toward the axis of the pentamer, has two anchorings: Arg85(n) and Asp88(n-1), belonging to the (n) and (n-1) pre-β5 strands respectively. In all CBX-bound structures (Sauguet *et al*., 2013, Fourati *et al*., 2020), each Arg77(n) side-chain has pivoted toward the periphery of the pentamer, and coordinates the CBX molecule bound in the inter-SU pocket(n). The side chain of Arg85(n) now interacts within the intra-SU pocket with Glu104(n) side-chain. And on the axial side of the (n) to (n+1) space/interface between two subunits, the Asp88(n) to Arg77(n+1) ion bridge is lost (Fig. 6*A*): the side-chains of the pre-β5 Asp86(n) and Asp88(n) residues point differently in the vestibule lumen. In all GLIC-CBX co-structures, the five inter-SU pockets are occupied, and the scheme occurs five times: the Asp88(n-1) to Arg77(n) bridge is lost, etc.

Therefore, binding of a CBX molecule at the inter-SU pocket (n) [and its coordination by Arg77(n)], occurs with, or requires, rupture of the Asp88(n-1)-Arg77(n) ion bridge, [and rupture of the Arg85(n)-Arg77(n) intra-SU interaction]. It may be that inter-SU CBX binding is facilitated when one of Arg77 anchorings (Asp88, Arg85) in its apo-GLIC orientation is missing. Consistently, positive modulation is favoured in the pre-β5 variant lacking Asp86 and Asp88.

### Release of two pre-**β**5 strand anchorings, and the pLGICs Cys-Loop (or Pro-Loop)

Conversely, CBX binding frees the pre-β5 strand(n) from both its Asp88(n)-Arg77(n+1) anchoring to the adjacent subunit, and its Arg85(n)-Arg77(n) intra-subunit anchoring. CBX binding also changes Arg85(n) orientation within the intra-SU pocket, ending in changing the pre-β5 strand orientation. We suggest that the pre-β5 strand motion is a major point in coupling CBX binding and pore gating. Given the peculiar properties of the Asp86-Asp88 pair of residues, we chose to use a double mutant to characterize a pre-β5 involvement. And we have not planned to further analyse the respective contributions of Asp86 *vs* Asp88 to the amazing properties described for the D86A-D88A variant.

In GLIC structures, the pre-β5 strand N-ter to C-ter orientation is “ascending”, *i*. *e*. toward the extracellular *apex* of the pentamer. The pre-β5 and β5 strand is adjacent and antiparallel (“beta sandwich”) to the “descending” β6 strand, which itself is ending in the β6-β7 Loop, homologous to the “Pro-Loop/Cys-Loop” known to be essential to gating in Eukaryote pLGICs. It should be emphasized here that Tyr102 and Glu104, analysed in our work as intra-SU pocket residues, belong to this β6 strand descending to the Pro-Loop/Cys-Loop. The general location in Eukaryote pLGICs of the pre-β5-β5 strand (or β4-β5 strand) forming Loop Ω, attached to the β5-β6 “sandwich” ending with the Pro-Loop/Cys-Loop), suggests that our mechanistic model, with inter-SU binding, and vestibular control of a pre-β5-β5 strand motion also dragging the Pro-Loop/Cys-Loop, may have some relevance to the mechanism of gating control by neurotransmitters in human pLGICs.

### Relevance of the model to Eukaryote pLGICs

Some authors questioned the presence in the ECD of Eukaryote pLGICs of a cavity homologous to the Prokaryote vestibular pocket, as this may end up in a new allotopic target site for NAM or PAM therapeutic compounds. Is it possible to awake a functional vestibular site in human pLGICs? The presence in type 3 serotonin receptors (5HT_3_Rs) of an empty cavity at this location was noticed by Hu *et al*. (2018), who also defined the β4-β5 loop as Loop Ω. And the point was analyzed systematically in Eukaryote pLGICs by Brams *et al*. (2020). They concluded that, as in 5HT_3A_ receptors, a vestibule cavity is accessible in several cationic pLGIC subunits, among which muscular type nicotinic receptor subunits, due to a “Ω-open” conformation of the loop. In other cationic pLGICs subunits however, including (non-α7) neuronal nicotinic subunits, the vestibule site entrance is obstructed by its own Loop Ω in a “Ω-in” conformation, and therefore non-accessible to ligands. Brams *et al*. also showed that MTSEA-biotin modification of engineered Cys mutants in the vestibule site (in particular Leu151 in the β6 strand) produces a PAM effect on the serotonin-activated 5HT_3A_ receptor, showing that it is possible to target the vestibule site for modulation of the 5HT_3A_ receptor, and putatively other Eukaryote pLGICs.

In anionic pLGICs, Loop Ω (“Ω out”, Brams *et al*. 2020) protrudes from subunit (n) (complementary side) into a cavity of the neighbour (n+1) subunit (principal side), on the axial/vestibule/lumen side of the ECD. This cavity (n+1), occupied by the neighbour subunit Loop Ω (n), corresponds to the Prokaryote vestibular pocket, and is therefore adjacent to the orthotopic, neurotransmitter binding site. In GLIC, FUMAR or CROTON binding, associated with Arg77 pivot rotation to the CBX-bound orthotopic/inter-SU site, releases the Asp88(n)-Arg77(n+1) ion bridge [the pre-β5 strand (n)-(n+1) anchoring], and frees Arg85 (the pre-β5 strand) from its intra-SU anchoring through other intra-SU residues. It may be hypothetized that the Asp88(n)-Arg77(n+1) ion bridge evolved toward the insertion of a whole protruding loop in a pocket within the neighbour subunit. It may be that the ancestral 2-arm system controlling the position of the pre-β5 strand (and β5-β6 sandwich, and Pro-Loop) from the orthotopic site in GLIC was replaced during evolution by a modern “joystick” (Loop Ω), hold in hand by a vestibular pocket, itself manipulated by the adjacent neurotransmitter binding site, a “joystick” which drags the β5-β6 sandwich ending with the Cys-Loop.

## Abbreviations list

AA: D86A-D88A pre-β5 GLIC variant, or D86A-D88A double mutation
ACET: acetic acid/acetate
BAPTA: 1,2-bis(*o*-aminophenoxy)ethane-*N*,*N*,*N’*,*N’*-tetraacetic acid
BUTYR: butyric acid/butyrate
CAFFE: caffeic acid/caffeate
CBX: carboxylic acid/carboxylate
CROTON: crotonic acid/crotonate
ECD: extracellular domain
FUMAR: fumaric acid/fumarate^-^/fumarate^2-^
GABA: *gamma*-aminobutyric acid
GLIC: *Gloeobacter violaceus* ligand-gated ion channel
GoF: gain of function
HEPES: *N*-(2-hydroxyethyl)piperazine-*N*′-(2-ethanesulfonic acid
intra-SU: intra-subunit
inter-SU: inter-subunit
LoF: loss of function
MES: 2-(*N*-morpholino)ethanesulfonic acid
mM: mmol/L
NAM: negative modulator of the allosteric transitions
PDB: Protein Data Bank
pHo: pH of the extracellular solution
pLGIC: pentameric ligand-gated ion channel
PAM: positive modulator of the allosteric transitions
PROPION: propionic acid/propionate
SUCCIN: succinic acid^-^/succinate^-^/succinate^2-^
TMD: transmembrane domain
VALER: valeric acid/valerate.

## Additional information

### Data Availability Statement

For patch-clamp electrophysiology data, representative raw current traces (current recorded *versus* time) are available in most of the Figures (Figs. 1, 2, 3, 4, 5, 7, 8; not in Fig. 6). For electrophysiology data presented as bar graphs, individual value data points (one per cell in each condition) are superimposed to mean ± SD (Figs. 3*B*, 4*B*, 5*CD*, 6*B*, 7*BC*, 8*BC*), and the number of cells tested is indicated in brackets in each bar graph category name. Mean ± SD is represented in Graphs for [concentration] – [current] “dose-response” curves (Fig. 2*BD*). The number of cells included is then indicated in the Figure Legend and/or Text. In all cases, “n” includes cells from at least two DNA transfections (tk-ts13 cells) or two DNA injections (*X*. oocytes). A statistical comparison of pairs of samples was performed in experiments with carboxylates on GLIC mutants (Figs. 4*B, 5CD*, 6*B*, 7*BC*, 8*BC*). P values are then indicated near bar graphs in the Figures, and most of the time also in Text.

## Author contributions

CVR set up patch-clamp, prepared some mutants used in Figs. 7&8, designed and performed patch-clamp experiments (including cell preparation), analysed data (including statistical treatment), prepared patch-clamp Figures, and performed part of *Xenopu*s oocyte electrophysiology. ÁN performed part of *Xenopus* oocyte electrophysiology and prepared Fig. 6A. KM performed molecular biology constructs and production. P-JC initiated and supervised the work. CVR wrote the first version of the article. All authors contributed to revising the article.

All authors have approved the final version of the manuscript and agree to be accountable for all aspects of the work. All persons designated as authors qualify for authorship, and all those who qualify for authorship are listed.

## Funding

ÁN was supported by the Agence nationale de la recherche (ANR grant 13-BSV8-0020 “Pentagate”). CVR and P-JC have permanent CNRS positions, KM has a permanent Institut Pasteur position. P-JC’s laboratory belongs to IP, CNRS and UPC, supported by public funding and donators.

### Ackowledgements

CVR wishes to thank Chantal Jan, Colette Jean, Corinne Balayira and Jamal Boutchlhit for technical assistance with lab-ware; Marc Gielen for building the oocyte set-up used for this report; Ákos Nemecz and Laurie Peverini for tutorials with molecular biology; Marc Delarue, Ludovic Sauguet, Zaineb Fourati-Kamoun for discussion; and Julien Lalande for reading the manuscript.

## First author Profile

CVR started electrophysiology in 1981, after listening to lectures by Philippe Ascher. She worked on various topics, including calcium-release activated calcium influx in excitable cells (Van Renterghem & Lazdunski 1994), and voltage-gated Na^+^ and Ca^2+^ channels. She enjoys *live* recording at the patch-clamp set-up. While considering proteins and ion channels, she develops more and more astonishment and admiration regarding the splendour of the living world. She is presently interested in voltage-dependent properties of pentameric ligand-gated ion channels.

## Authors’ Translational Perspective (*not provided*)

